# ROS-Degradable Polythioketal Urethane Foam Dressings to Promote Porcine Skin Wound Repair

**DOI:** 10.1101/2021.05.21.445175

**Authors:** Prarthana Patil, Katherine A. Russo, Joshua T. McCune, Alonda C. Pollins, Matthew A. Cottam, Bryan R. Dollinger, Juan M. Colazo, Fang Yu, John R. Martin, Mukesh K. Gupta, Nancy L. Cardwell, Jeffrey M. Davidson, Callie M. Thompson, Adrian Barbul, Alyssa M. Hasty, Scott A. Guelcher, Craig L. Duvall

## Abstract

Impaired skin healing and progression into chronic wounds is a prevalent and growing medical problem. Porous, resorbable biomaterials can be used as temporary substrates placed into skin defects to support cell infiltration, neo-tissue formation, and remodeling of nonhealing wounds. Naturally-derived biomaterials have promising healing benefits, but their low mechanical properties and exuberant costs limit their performance and use. Synthetic materials can be affordably manufactured and tuned across a broader range of physiochemical properties, but opportunities remain for tailoring them for ideal host immune and regenerative responses. Polyesters are the most clinically-tested class of synthetic biomaterials, but their hydrolysis releases acidic degradation products that can cause autocatalytic degradation processes that are poorly controlled and are not tied to cellular or other biologic activities. Here, we systemically explored a series of ROS-degradable polythioketal (PTK) urethane (UR) foams with varied hydrophilicity as an alternative class of synthetic biomaterials for wound healing. It was found that the most hydrophilic PTK- UR variant, which had 7 ethylene glycol (EG7) repeats flanking each side of each thioketal bond, had the highest ROS reactivity of the PTK-URs tested. In an in vivo porcine excisional skin wound healing model, hydrophilic EG7 PTK-UR foams more effectively promoted tissue integration, ECM deposition, and re-epithelialization of full-thickness skin wound compared to more hydrophobic PTK-UR variants. Resolution of type 1 inflammation and lower foreign body response to scaffold remnants was also observed for EG7 versus more hydrophobic PTK-UR scaffolds. Finally, porcine wound healing studies showed that EG7 PTK-UR foams had similar wound healing response to a collagen-based clinical gold standard product, Integra Bilayer Wound Matrix (BWM), while outperforming polyester UR foam-based NovoSorb Biodegradable Temporizing Matrix (BTM) with respect to increased ECM production, vascularization, and biomaterial-associated immune phenotype. In sum, PTK-UR foams warrant further development toward a new class of synthetic biomaterial foams for skin wound healing applications.

## Introduction

Chronic wounds affect approximately 4.5 million Americans and cause a large economic burden in the United States, with costs for wound management alone estimated to be as high as $96.8 billion per year (*1, 2*). Diabetes, obesity, and cardiovascular disease predispose patients to incomplete wound closure, often requiring surgical intervention and in severe cases, lower limb amputation (*1, 3, 4*). Chronic ulcers take approximately 1 year to heal on average in adults; recurrence rate is high, and 5-year mortality rate is ∼40% for patients who experience diabetic skin ulcers (*5, 6*). Chronic wounds are characterized by persistent, low-grade inflammation with an influx of polymorphic mononuclear leukocytes (PMNs) that secrete inflammatory cytokines, proteolytic enzymes that breakdown extracellular matrix (ECM) and growth factors, and reactive oxygen species (ROS) (*7*). Excess ROS causes oxidative stress and reduces angiogenesis, ECM deposition, and healing capacity of fibroblasts and keratinocytes (*8*). Unclosed wounds also have a high incidence of bacterial infection and biofilm formation, which further exacerbate the local inflammatory imbalance (*9*).

Standard of care for treating chronic wounds includes tissue debridement, moisture control, and use of biomaterials with or without additives to reduce infection. Biomaterials, primarily comprising either naturally-derived or synthetic polymers, can be used as artificial dermal substitutes that are placed permanently into the wound bed where they act as a resorbable, provisional matrix that provides mechanical support and promote cellular attachment, ECM deposition, and consequently accelerated defect repair (*10*). Acellular wound dressings based on animal-derived ECM components are particularly well-established. The first of its kind, Integra Bilayer Wound Matrix (BWM), designed by Yannes et al (*11*), has a long clinical track record; in diabetic foot ulcers, Integra BWM achieves full closure in 51% of wounds by 16 weeks, compared to 32% by standard wound care (*12*). Oasis wound dressings manufactured from porcine subintestinal submucosa (SIS) is another popular option that decreases time required for complete wound closure and reoccurrence of venous ulcers (*13*). Many ECM-based wound dressings have been developed for clinical use (*14*), but they suffer from challenges including reliable raw material sourcing, reproducible processing techniques, cross-species immunogenicity risk, and exuberant costs (*15*).

Fully synthetic dermal materials are a promising alternative that provides more reproducible and cost-effective product fabrication, while also affording greater potential for tuning of chemical composition, microarchitecture, physiochemical properties, and degradation profiles. Hydrolytically-degradable polyesters (PE) are the main class of synthetic polymers that have been developed into clinically-approved, resorbable wound dressings (*16*). Two primary examples are Restrata^®^ and NovoSorb^®^ biodegradable temporizing matrix (BTM). Restrata^®^ is an electrospun PLGA (10:90) and polydioxanone mat, while NovoSorb^®^ is a polyurethane foam made from ethyl lysine diisocyanate, lactic acid/ethylene glycol chain extender, and PCL1000 polyol with a removable polyurethane overlayer (*17, 18*). Restrata^®^, and NovoSorb^®^ have both gained broad FDA approval for indications spanning traumatic and post-surgical wounds, burns, pressure ulcers, and diabetic ulcers. Highly porous polyurethane foams are advantageous because they reduce the required production materials, quantity of degradation products, and production cost. Polyurethanes are also a tunable chemistry with properties that can be controlled through variation of either the polyol or isocyanate-containing component; isocyanates have been paired with aliphatic PE such as poly(lactic acid) (PLA) (*19*), poly (lactic acid -glycolic acid) (PLGA), polycaprolactone (PCL) (*20*), and their blends to address tissue engineering needs in bone regeneration (*21*), soft tissue repair (*22*) and drug delivery (*23, 24*). PE based materials do, however, suffer from the release of acidic degradation products which can activate an autocatalytic degradation profile (*25*) that is unfavorable in the context of mechanical strength and biomaterial associated inflammation (*26*).

A newer class of stimuli responsive materials, polythioketals (PTKs), are hydrolytically stable, however, undergo oxidative degradation in the presence of cell-derived ROS cues, without the formation of inflammatory/acidic by-products. We have previously shown that PTK-based polyurethane scaffolds provide more temporally-controlled degradation kinetics compared to PE materials and promote robust tissue integration in rat and porcine skin wounds (*27, 28*). In addition to PTK-UR scaffolds, we have shown that PTK crosslinked PEG-MAL hydrogels promote tissue infiltration and vascularization in mice, confirming the biocompatibility of hydrophilic PTK-based materials in physiologically relevant environments (*29*).

Here, we posited that relative scaffold hydrophilicity was an important parameter in optimization of wound healing performance of PTK urethane (PTK-UR) dermal substitutes. This hypothesis was motivated by studies showing that integration of PEG can increase degradation rate and tissue infiltration of PE-UR scaffolds in vivo (*19, 24*) and observations that zwitterionic polymers can decrease fibrosis and trigger a regenerative immunophenotype (*30*). Toward this end, we generated a library of ROS-responsive PTK-UR foams with controlled variation of the PTK diol component to have different quantities of ethylene glycol (EG) units between the TK bonds in the polymer backbone. We hypothesized that increased hydrophilicity of PTK-UR scaffolds would increase thioketal group reactivity with water soluble ROS, thus improving scaffold ROS scavenging/degradation ultimately promoting faster and more favorable rate of material resorption in vivo. The primary goals of these studies were to establish the functional importance of PTK- UR scaffold hydrophilicity on wound healing and to benchmark leading PTK-UR formulations against current standards of care in a clinically-relevant porcine skin wound repair model.

## Materials and Methods

### Study design

The current studies tested a library of ROS degradable PTK-UR scaffolds in a porcine wound model. All surgical procedures were reviewed and approved by Vanderbilt University’s Institution Animal Care and Use Committee. We created twenty-eight, 2 x 1 cm full thickness wounds on the dorsal region of healthy adolescent Yorkshire female pigs. An adequate distance (≥ 2 cm) was maintained between each wound to avoid overlapping treatment effects while simultaneously enabling higher throughput screening of multiple biomaterial formulations, (n=3/4 per treatment in each animals), as previously described (*28, 31, 32*). Treatment placement was randomized throughout the dorsal region to avoid anatomical bias. To evaluate host response and early to mid-stage immune response to the implanted scaffolds, we euthanized at 10 days post scaffold implantation and harvested tissue for downstream analysis. Full-thickness biopsies from within the wound margins were collected post anesthesia, prior to euthanasia. To further evaluate rate of wound closure and resurfacing of PU treated wounds (n=6-10), animals were euthanized 24 days post wounding, and tissue was harvested for histological and gene expression analyses. In a separate experiment, we treated porcine full thickness wounds (n=6-9) with our lead PTK-UR formulation, Integra BWM, and NovoSorb BTM. In this study, we evaluated wound closure and blood perfusion over time, in addition to gene expression and histological measurements at the endpoint at 31 days post wounding. Dressing regime was consistent across treatments and animals and changed three times a week. Animals received weekly antibiotic injections to address potential bacterial infections along with analgesics to manage pain. Pre- mature loss of scaffolds from the wounds due to mechanical disturbance by the animal prior to the pre- determined end point was a criterion for exclusion of that sample from outcome analysis. None of the treatments resulted in animal weight loss during the course of the experiment. All experiments in vitro were repeated three times. Pathohistological scoring was done in a treatment-blinded fashion.

### Synthesis of EG0(BDT), EG1(MEE), and EG2(EDDT) PTK diols

EG0, EG1 and EG2 PTK diols were synthesized as previously publish by Martin et al (*27, 29*). 1,4 butanedithiol (BDT, EG0), mercaptoethyl ether (MEE, EG1), and 2,2’-(ethylenedioxy) diethanethiol (EDDT, EG2) were synthesized through condensation polymerization with dimethoxy propane (DMP) using *p*-toluene sulphonic acid (pTSA) as a catalyst under inert conditions (Figure S1). pTSA was dissolved in concentrated hydrochloric acid (HCl) and heated to 50°C. The acid was allowed to cool, and pTSA crystals were collected, rinsed with cold HCl, and vacuum dried for 2 h. A tri-neck flask attached with an addition funnel was charged with anhydrous acetonitrile and BDT/MEE/EDDT (1 mol eq, 0.400 mol) stirred continuously and heated to 80°C followed by the addition of re-crystallized pTSA (0.005 mol eq, 0.002 mol). DMP (0.83 mol eq, 0.334 mol) dissolved in ACN was added dropwise to the monomer mixture through the addition funnel. The reaction was allowed to stir overnight after which the crude polymer was concentrated under rotary evaporation, precipitated 3X in cold ethanol and vacuum dried. To functionalize the resulting dithiol polymers, PTK dithiol polymers (1 mol eq, 0.060 mol) were dissolved in anhydrous THF followed by the addition of CsCO3 (5 mol eq, 0.15 mol). 2-bromoethanol (5 mol eq, 0.15 mol) was added dropwise to the reaction and allowed to proceed for 4 hours. The resulting polymer was filtered, concentrated, precipitated 3X in ethanol, and vacuum dried. Polymer composition was evaluated using ^1^H nuclear magnetic resonance (NMR) spectroscopy. Respective PTK diol polymers were dissolved in deuterated chloroform (CDCl3) and analyzed at 400 Hz ^1^H NMR; chemical shifts were reported as δ values in ppm relative to deuterated CDCl3 (δ = 7.26). BDT PTK diol; δ=1.60 (s, 6H), δ=1.65-1.69 (m, 4H), δ=2.58-2.61 (t, 4H), δ=2.69-2.73 (m, 2H), δ=3.68-3.72 (m, 2H). MEE PTK diol; δ=1.60 (s, 6H), δ=2.72- 2.76 (m, 2H), δ=2.78-2.82 (t, 4H), δ=3.58-3.64 (m, 4H), δ=3.68-3.74 (m, 2H). EDDT PTK diol; δ=1.60 (s, 6H), δ=2.72-2.74 (m, 2H), δ=2.77-2.80 (t, 4H), δ=3.56-3.60 (m, 8H), δ=3.68-3.74 (m, 2H)

### Synthesis of EG7 PTK diols

Synthesis of thioketal diethylamine was adapted from previously published work as shown in Figure S2 (*33*). Cysteamine (1 mol eq, 0.47 mol) was dissolved in anhydrous methanol under nitrogen flow, followed by the addition of triethylamine (1.5 mol eq, 0.70 mol). The reaction was stirred on ice for 15 mins. Ethyl trifluoroacetate (1.2 mol eq, 0.56 mol) was added to the solution dropwise and allowed to stir on ice for 1 hour. The reaction mixture was brought to room temperature and stirred overnight. The reaction mixture was concentrated on a rotary evaporator before it was dissolved in water and extracted into ethyl acetate (3X). The combined organic layer was dried over anhydrous magnesium sulfate and concentrated in vacuo. The crude product was further purified by silica gel chromatography using a gradient of 100% to 70:30 ethyl acetate:hexane. The product (compound [1], Figure S2), a colorless liquid, was analyzed using 400Hz ^1^H NMR; chemical shifts were reported as δ values in ppm relative to deuterated CDCl3 (δ = 7.26) as follows; δ=2.7-2.76 (m, 2H), δ=3.52-3.56 (m, 2H), δ=7.01 (s, 1H) Compound [1] (1 mol eq, 0.42 mol) was dissolved in ACN and charged with N2 gas followed by the addition of BiCl3 (0.025 mol eq., 0.005 mol) as a catalyst. Using an addition funnel, DMP (1 mol eq., 0.192 mol) dissolved in ACN was added to the flask dropwise. The reaction was then allowed to stir at room temperature for 24 hours. Acetonitrile was removed from the mixture via rotary evaporation, and the crude product was filtered through silica in a mixture of 70-30 hexane/ethyl acetate. The flow through was collected and concentrated in vacuo. The product (compound [2], Figure S2) (65% yield), a white solid, was analyzed through 400Hz ^1^H NMR in CDCl3. ^1^H NMR chemical shifts were reported as δ values in ppm relative to deuterated CDCl3 (δ = 7.26) as follows; δ=1.60 (s, 6H), δ=2.8-2.85 (t, 4H), δ=3.56-3.61 (m, 4H), δ=6.8 (s, 2H)

Compound [2] (129 mol) was dissolved in minimal volume of methanol (MeOH), and a saponification reaction was carried out in an aqueous 6M NaOH solution. The mixture was allowed to stir for 4 hours at room temperature. The product (compound [3]) was extracted into dichloromethane (DCM, 3X). The combined organic fractions were dried using anhydrous magnesium sulfate and concentrated under vacuum to obtain the product (60% yield), a pale-yellow colored oil. Product was analyzed through 400Hz ^1^H NMR in DMSO (δ = 2.5) as follows; δ=1.53 (s, 6H), δ=2.58-2.62 (t, 4H), δ=2.66-2,69 (m, 4H) Hydroxyl activated PEG 400 was synthesized as previously described (*34*). PEG 400 (Mn = 400 Da) was dried at 80°C under vacuum for 24 hours. Dried PEG (1 mol eq, 0.10 mol) was dissolved in anhydrous DCM along with triethylamine (1.2 mol eq., 0.12 mol). In a separate round bottom flask, p-nitrophenyl chloroformate (1.1 mol eq., 0.110 mol) was dissolved in anhydrous DCM and slowly added dropwise to the PEG400/TEA mixture over ice. The reaction was allowed to stir for 30 mins on ice, followed by 24 hours at room temperature. Tetraammonium hydrochloride salts that were formed during the reaction were removed via filtration and the DCM was removed on a rotary evaporator. The product was concentrated to obtain monofunctionalized PEG 400 nitrophenyl carbonate (PEG400-mNPC). This step also yielded a small fraction of homobifunctional PEG 400 dinitrophenylcarbonate (NPC-PEG400-NPC). Product was analyzed through 400Hz ^1^H NMR in DMSO (δ = 2.5) as follows; δ=3.5-3.6 (m, 28H), δ=3.71 (m, 2H), δ=4.4 (m, 2H), δ=7.6 (d, 2H), δ=8.3 (d, 2H). To synthesize EG7 PTK diol, hydroxyl activated PEG nitrophenyl carbonate (2.4 mol eq, 0.034 mol) was dissolved in anhydrous DCM. A mixture of the thioketal ethyldiamine crosslinker (1 mol eq, 0.014 mol) and triethylamine (2.4 mol eq, 0.034 mol) was dissolved in DCM, added dropwise to the PEG400-mNPC/NPC-PEG400-NPC solution, and allowed to react for 4 hours at room temperature. The crude mixture was then precipitated 3X in cold diethyl ether to remove p-nitrophenol impurities, and additional purification was carried out through silica column chromatography. Crude product was loaded onto a silica packed column, and fractions of clean product were eluted using a mixture of DCM:MeOH at gradients of 0 to 10% MeOH. The fractions were then dried in vacuo and the product (50% yield) was analyzed through 400Hz ^1^H NMR in DMSO (δ = 2.5) as follows; δ=1.54 (s, 12H), δ=2.58-2.62 (t, 8H), δ=3.11-3.16 (m, 8H), δ=3.49-3.53 (m, 94H), δ=3.61 (t, 8H), δ=4.17 (s, 4H), δ=7.34 (s, 4H).

### Polyester polyol synthesis

Trifunctional polyester polyols (900t and 1500t) were synthesized as previously published(*35*). Briefly, glycerol was dried for 48 hours at 80°C under vacuum. Dried glycerol was added to a tri-neck flask, charged with N2 followed by the addition of 60% ε-caprolactone (CL), 30% glycolide (GA), and 10% D, L-lactide (LA) along with a stannous octoate catalyst to yield a 900Da triol and 1500Da triol.

### Determination of molecular weight and hydroxyl number

Polymer molecular weight (Mn) and polydispersity were determined through dimethylformamide (DMF) gel permeation chromatography (GPC) with inline RI and light scattering detectors, based on dn/dc values obtained from an offline refractometer. To analyze the OH number of the resultant polymeric diols, ^19^F NMR spectroscopy was used as adapted from Moghimi et al (*36*). 0.03 mmol of PTK diol was dissolved in deuterated acetone. 0.1 wt% (dibutyltin dilaurate) DBTDL as catalyst and 0.081 mmol of α, α, α-trifluorotoluene (CF3-Ph) as internal standard was added to the PTK diol solutions. Finally, 0.3 mmol of reagent 4-fluorophenyl isocyanate (4-F-Ph-NCO) was mixed in with the reactants and allowed to react for 15 mins at room temperature. The ^19^F NMR spectrum in deuterated acetone was recorded as follows; δ= - 63.23 (s, 3F, CF3-Ph), δ= -117.6 (m, 1F, 4-F-Ph-NCO), δ= -122.2 (s, 2F, 4-F-Ph-NH-CO-O-PTK).

Theoretical hydroxyl number of each PTK diol was calculated by:

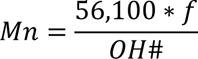

where 56,100 represents the molecular weight of KOH in mg/mol, *f* represents the hydroxyl functionality of the PTK (2 for a linear homobifunctional PTK polymer) and Mn represents the number-average of the PTK polymer

### Fourier Transform Infrared Spectroscopy (FTIR) and Differential Scanning Calorimetry (DSC)

10 uL of each PTK diol were applied in a thin layer between two 25 x 4 mm KBr glass windows and loaded into a Bruker Tensor 27 FTIR Spectrometer. Absorbance at 3400 cm^-1^ was used to identify OH groups for each PTK diol. In addition, absorbance peaks at 3330 cm−1 (N–H stretching) and a sharp band at 1630 cm−1 (carbonyl groups in urea bonds C=O) was used to confirm the presence of NHCOO bonds in EG7 PTK diols.

For glass transition temperature (Tg) analysis using DSC, samples ranging in mass from 5 to 9 mg were heated from -80.0°C to 20.0°C at a rate of 10°C min^-1^. Samples were then cooled to -80.0°C at a rate of 10°C min^-1^, and heated a second time to 30°C at a rate of 10°C min^-1^. All transitions were obtained from the second heating run.

### Urethane scaffold fabrication

The PTK and PE urethane (UR) scaffolds were fabricated using liquid reactive molding. PTK-diols were mixed with calcium stearate (pore opener), TEGOAMIN33 (catalyst), water (blowing agent), and sulfated castor oil (pore stabilizer) for 30 seconds at 3300 rpm in a Speed Mixer Lysine triisocyanate (LTI) was added and mixed for an additional 30 s at 3300 rpm. The mixture was allowed to rise, set, and harden overnight before the scaffolds were demolded and trimmed to shape. The target isocyanate: hydroxyl (NCO: OH) was set to 115:100 where the OH equivalents is obtained from polyol’s molecular weight and OH# obtained from ^19^F NMR. The mass of each component for each scaffold formulation denoted as parts per hundred-part polyol (PPHP) was individually optimized and is outlined in Table 1 (supplementary figures).

**Table 1:**
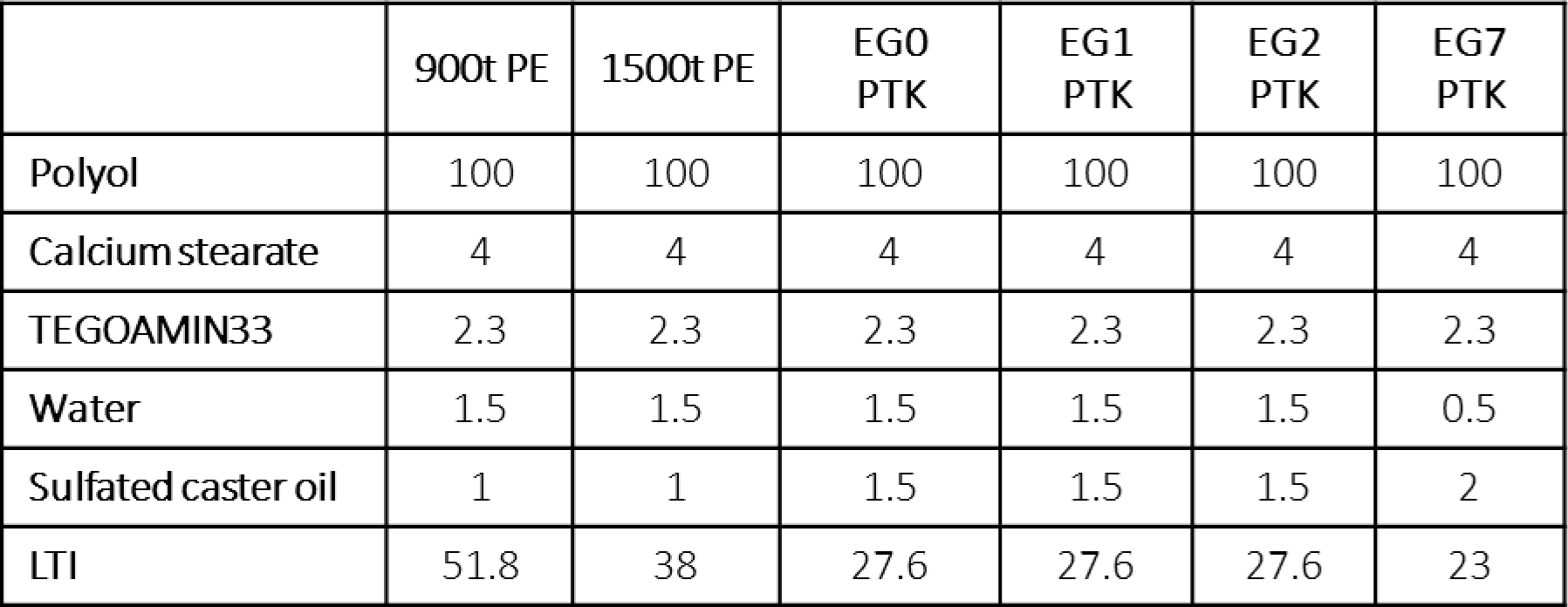
PTK and PE-UR scaffold fabrication components.

|  | 900t PE | 1500t PE | EG0<br>PTK | EG1<br>PTK | EG2<br>PTK | EG7<br>PTK |
| --- | --- | --- | --- | --- | --- | --- |
| Polyol | 100 | 100 | 100 | 100 | 100 | 100 |
| Calcium stearate | 4 | 4 | 4 | 4 | 4 | 4 |
| TEGOAMIN33 | 2.3 | 2.3 | 2.3 | 2.3 | 2.3 | 2.3 |
| Water | 1.5 | 1.5 | 1.5 | 1.5 | 1.5 | 0.5 |
| Sulfated castor oil | 1 | 1 | 1.5 | 1.5 | 1.5 | 2 |
| LTI | 51.8 | 38 | 27.6 | 27.6 | 27.6 | 23 |

### Urethane film synthesis and contact angle measurement

PE-UR and PTK-UR films were synthesized by combining polymeric diols and bismuth neodecanoate bismuth (a gelling catalyst) in a 22 mm polyethylene mold (Fisher Brand) for 30s at 3300 rpm. LTI was then added to this mixture and mixed for additional 60s. Urethane films were allowed to cure for 24 hours at RT before being demolded and punched into 4 mm circles using a biopsy bunch.

Contact angle measurements were carried out using a goniometer. A 4µl water droplet was dispensed onto the surface of the urethane film and allowed to equilibrate for 10 mins. The resulting contact angle formed between the water droplet with the film was measured using ImageJ. Bilateral contact angle measurements were obtained for each film, and averaged values across three different batches of urethane films were recorded for each formulation.

### Sol fraction

Fabricated scaffolds were weighed to obtain initial dry mass (*Mo*) and incubated in DCM for 24 hours. Scaffolds were then removed from DCM, air dried and weighed to obtain the final mass (*Mf*). Soluble fraction (*fs*) was calculated using the following equation:

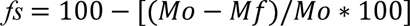

### Swell ratio

To determine the swelling ratio of PTK-UR and PE-UR scaffolds, 10 mg (dry mass, *Md*) scaffolds were incubated in PBS overnight and weighed to obtain swollen mass (*Ms*). The swell ratio was calculated using the following equation:

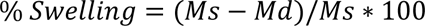

### Dynamic mechanical analysis

Mechanical properties of PTK-UR and PE-UR scaffolds were measured in compression using the TA Q800 Dynamic Mechanical Analyzer. 6 x 6 mm cylindrical scaffold punches were hydrated in PBS for 24 hours prior to measurement. Scaffolds were longitudinally compressed under a preload force of 0.1N at 10% strain min^-1^ until 60% strain was reached. The longitudinal compression was repeated three times and values averaged per sample. Young’s modulus for each sample was calculated from the linear region of the stress-strain curve after initial toe-in. A total of 3 scaffold pieces across three separate batches were averaged to obtain the Young’s modulus.

### In vitro degradation of PTK-UR scaffolds

Pre-soaked and dried 10 mg scaffold pieces (Mo) fabricated from PE and PTK diols were incubated in PBS and increasing concentrations of hydrogen peroxide/cobalt chloride (H_2_O_2_/CoCl_2_) solutions to generate hydroxyl radical; 0.2% H_2_O_2_/0.001M CoCl_2_, 2% H_2_O_2_/0.01M CoCl_2_, or 20% H_2_O_2_/0.1M CoCl_2_. Scaffolds were incubated in 2 mL of treatment media for up to 30 days with media replaced every other day. At pre- determined time points, scaffolds were removed from the treatment media, rinsed, dried, and weighed to obtain remaining mass (Mt) of the scaffold. Three independent scaffold pieces were tested at each time point. Degradation was recorded as percent mass loss for each time point using the following equation:

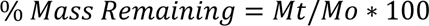

### Mathematical modeling of PTK-UR scaffold degradation

MATLAB was used to fit the oxidative degradation kinetics of PTK-UR scaffolds to a first order mathematical model based on H_2_O_2_ concentration using

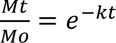

where *Mt* represents the final mass of scaffold time at time *t*, *Mo* represent*s* the initial scaffold mass, and *k* represents the degradation rate constant. Non-linear regression was used to fit the experimentally obtained scaffold degradation data to the first order mathematical model to determine the degradation rate constant for each scaffold with respect to the degradation media concentration.

### Scanning electron microscopy of urethane scaffolds

Scaffold pieces (60-80 mg) fabricated using PE and PTK diols were incubated in PBS or 2% H_2_O_2_ with 0.01M CoCl_2_ for up to 15 days. At each time point, scaffolds were removed from their respective treatments and washed in a series of graded ethanol (30%, 50%, 70%, 80%,90%, 95% and 100%) for 10 mins to remove the aqueous content. Scaffolds were then dried via critical point drying and weighed to obtain final scaffold mass. Scaffold pieces were sputter coated in gold using a Cressington Sputter Coater. Scaffolds were imaged using Everhart-Thornley secondary electron detector at a beam voltage of 3.00 keV or 10 keV. Pore size was quantified through ImageJ using SEM images.

### DPPH radical scavenging assay

The radical scavenging potential of PE-UR and PTK-UR scaffolds was assessed through using 1,1- diphenyl-2-picrylhydrazyl (DPPH) assay (*37*). DPPH was dissolved in a mixture of 80:20 ethanol/H2O (v/v%) at 200 µM concentration. Pre-fabricated PE and PTK urethane scaffolds were trimmed to 10 mg pieces (n=3), treated with 2 mL of DPPH solution, and incubated at 37°C on an orbital shaker for 36 hours. 100 µL of incubation solution was sampled at 1, 2, 3, 6, 12, 24 and 36 h time points and the absorbance (Asample) was measured at 517 nm and compared to control DPPH solution absorbance (Acontrol). The scavenging potential is reported as following

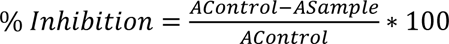

### ROS cell protection assay

Cell protection against cytotoxic level of oxidative stress was tested in NIH 3T3 fibroblasts. Briefly 10,000 cells were seeded in a 96 well plate and allowed to adhere overnight. Cells were sequentially treated with 100 µL of either 25 µM H_2_O_2_ or 50 µM H_2_O_2_ and 100 µL of polyol solution. PTK diol mass for each treatment was adjusted to match the overall thioketal molar content for each group. PE polyol controls were formulated at 0.5 mg/mL to match the highest PTK diol mass. 24 hours post treatment; cell viability was detected using CellTitre-Glo Luminescent Cell Viability Assay (Promega) and measured on IVIS 200. Signal from treatment groups was normalized to untreated cells and expressed as relative cell viability.

### Porcine excisional wound model and PTK-UR scaffold screening

All surgical procedures were reviewed and approved by Vanderbilt University’s Institution Animal Care and Use Committee. Adolescent female Yorkshire pigs were anesthetized with a cocktail of telezol (4.4mg/kg), ketamine (2.2mg/kg), and xylazine (2.2mg/kg) administered intramuscularly and maintained under isoflurane for the duration of the surgery. Shaved dorsal skin was sterilized using 70% ethanol, chlorohexidine, and betadine washes. Twenty-eight, full thickness 2 x 1 cm excisional wounds were created on the dorsal region of the pig. Ethylene oxide sterilized scaffold pieces were trimmed to 0.2 x 2 x 1cm and implanted into the wounds after hemostasis was achieved. Following implantation, wounds were covered with layerings of Mepilex Transfer (Molnlycke Healthcare, Sweden), Mextra absorbent pad (Molnlycke), Opsite adhesive (Smith&Nephew, St. Petersburg, FL), MediChoice Tubular Net Bandage (Owens & Minor, Mechanicsville, VA), and Vetwrap (3M, St. Paul, MN) bandaging over the wound. Following surgery, dressings were changed every 2-3 days. Antibiotics (Excede; Zoetis) were administered weekly during the period of the study. Buprenex was given immediately post-surgery, and analgesia was maintained for 7 days post-op using a transdermal Fentanyl patch (50 mcg/h) applied at the time of surgery and changed every 2-3 days. Pigs were euthanized at 10 and 24 days post-surgery; wound samples were collected using 2 mm biopsy punches prior to euthanasia for downstream gene expression analysis. Post euthanasia, wound tissue was excised for histological analyses.

### Testing of lead candidate against clinical controls

Integra Bilayer Wound Matrix (BWM) was doused in sterile saline, trimmed to 2 x 1 cm pieces, and placed matrix side down into the excisional wound. The protective silicone layer was removed 14 days post application, according to manufacturer guidance. NovoSorb Biodegradable Temporizing Matrix (BTM) scaffolds were trimmed to 2 x 1 cm pieces and applied matrix side down into the wound. Sealing polyurethane membrane was removed 21 days post application according to manufacture guidance. Treatments were randomized with regards to wound location to avoid any bias. The wounds were dressed and secured with additional bandaging as described above. Pigs were euthanized 31 days post-surgery; wound samples were collected for histology and gene expression analysis.

### Laser Doppler perfusion imaging

A laser Doppler imager (moorLDI2-HIR; Moor Instruments, Wilmington, DE) was used to track the perfusion within the wounds over the duration of the study. Flux images of wounds were obtained after scaffold implantation and subsequently during dressing changes at days 1, 4, 10, 17, 24, and 31 post-surgery. Blood perfusion within a 2 x 1 cm region of interest as well as uninjured skin was quantified using Moor analysis software. Data is presented as total flux within the wounds, normalized to the total flux recorded in normal skin.

### Histology and quantification

Explanted tissue samples were fixed in 10% neutral buffered formalin for 48 hours. Tissue samples were then dehydrated in a graded ethanol series, exposed to xylene, and embedded in paraffin. Paraffin blocks were sectioned at 5µm thickness and processed for histology and immunohistochemistry. Tissue sections were deparaffinized (xylene and ethanol gradients) and rehydrated in Tris-Buffered Saline, 0.1% Tween 20 (TBST) buffer. Masson’s trichrome staining and H&E (hematoxylin and eosin) staining were performed according to the manufacturer’s recommendation. Antigen retrieval was performed using citrate-based pH 6 antigen retrieval solution (Dako) for 1 min at 120°C and allowed to cool to 90°C in a decloaking chamber. Sections were then incubated for 40 mins in 3% H_2_O_2_ TBST solution and blocked with protein block (Dako) for 20 mins. Sections were further incubated in rabbit anti-von Willebrand factor (vWF) antibody, mouse anti-cytokeratin14 (CytoK14) antibody, mouse anti-αSMA antibody, rabbit anti-CCR7 antibody, rabbit anti-arginase-1 (Arg-1) antibody, rabbit anti-CD3 antibody, rabbit anti-myeloperoxidase (MPO) antibody, or rabbit anti-CD206 antibody for 60 mins at room temperature Secondary antibodies donkey anti-mouse HRP and donkey anti-rabbit HRP (Dako) were applied for 30 mins at RT and samples were exposed DAB substrate (Dako) for 5 mins. Slides were then rinsed in TBST buffer, dehydrated in graded ethanol and xylene solutions, and mounted with Acrytol mounting media.

Histological sections were imaged and evaluated using Metamorph Imaging Software and ImageJ to assess granulation tissue area, wound thickness occupied by scaffold, and tissue infiltration within the scaffold pores. Granulation tissue area was defined as cross-sectional area occupied by the scaffold and neo-tissue. Wound thickness occupied by scaffold was defined as the longitudinal thickness of the new granulation tissue occupied by the bulk scaffold/scaffold remnants. Tissue infiltration was defined as percent area occupied by new granulation tissue within scaffold pores averaged across 3 fields of view per sample. Brown pixels, positive immunoreactivity for diaminobenzidine (DAB), were quantified for each IHC marker (CytoK14, CD206, MPO, CD3 and vWF). Epithelialization was quantified as percent wound length covered by CytoK14 positive epidermal layer. vWF, MPO, CD206, and CD3 were quantified and expressed as percent ROI area occupied by positive brown pixels. A minimum of three fields of view per wound sample were analyzed and quantified. Analysis of CCR7 IHC was performed on color deconvolved (Image J Plugin) DAB-hemotoxylin images. Plot intensity measurement on DAB isolated images were performed up to 150 µm from the edge of the scaffold. Gray values (0–255) were inverted and background subtracted to obtain measurements for CCR7 staining intensity.

### Wound Scoring

Treatment blinded histomorphometry analysis of trichrome and H&E stained tissue sections was used for histological scoring of wounds based on a composite evaluation of five parameters: extent of granulation tissue formation (G), degree of vascularization (V), re-epithelialization (E), collagen deposition (amount and fiber orientation) (C), and inflammation (I). A serial section stained with trichrome stain was scored to estimate the extent of and organization of new connective tissue. Each parameter is rated on a 1-5 score, ranging from little or no activity to near-complete reorganization of tissue. The overall score comprising of G+V+E+C-(I/2) was assigned to each wound with a higher score indicating a more favorable wound healing response. Semi-quantitative analysis for the formation of foreign body giant cells (FBGC) was performed on wounds treated with EG7 PTK-UR, NovoSorb, and Integra. A FBGC score between 1-5 was assigned to each wound and subtracted from the total wound healing score to account for foreign body response to the implanted materials.

|  | **Neovascularization** | **Granulation Tissue Maturation** | **Epithelialization** | **Inflammation** | **Collagen** |
| --- | --- | --- | --- | --- | --- |
| **0** |  |  | No migration of epidermis from wound edges |  |  |
| **1** | Little evidence of endothelial cell ingrowth into wound site | Little evidence of mesenchymal cell infiltration; eschar predominant | Little migration of epidermis from wound edges over granulation tissue | Mostly resolved inflammatory response. No foreign body giant cells | Little or no evidence of collagen fibers. Provisional matrix consists of fibrin. |
| **2** | Angiogenic response at the wound margins and base; some extravasation | Active expansion of mesenchymal infiltration from the base and margins of the wound, accompanied by neutrophils and mononuclear cells | Initial progression of epidermal resurfacing with less than 50% coverage and hyperplasia | Residual inflammatory infiltrate with some giant cells and mild mononuclear cell infiltrate | Initial signs of collagen accumulation consisting of very fine fibrils in a random orientation. |
| **3** | > 50% of granulation tissue shows neovascularization | Continued filling of the wound site and displacement of the eschar above the wound surface | 50-90% coverage of wound site. Clear polarized differentiation of epidermis | Substantial inflammatory infiltrate dominated by mononuclear cells but significant persistence of polymorphonuclear cells, frequently near wound surface | Moderate amount of collagen deposition |
| **4** | Extensive vascularization with pruning and maturation of neovessels | Accumulation of extracellular matrix, reduced density of fibroblasts, and evidence of collagen fiber organization | 90-100% coverage of wound site with initiation of rete peg formation and advanced epidermal differentiation | Heavy inflammatory infiltrate resulting in impairment of healing progression, and invasion of adjacent unwounded tissue. | Loosely assembled parallel collagen fibers |
| **5** | Advanced vascular maturation with thinning of capillary networks | Extensive reconstruction of dermal architecture with absence of inflammatory infiltrate | Full keratinized epidermis with rete pegs and return of epidermal hyperplasia to thickness of adjacent, unwounded skin | Wound site dominated by inflammatory cells, including neutrophils, with consequent retardation of either granulation tissue or epidermal development. | Highly aligned collagen fiber networks with tendency to form reticular arrays. Reduced fibroblast density. |

### Gene expression analysis of granulation tissue

Wound samples were excised at pre-determined time points and stored in RNAlater at -20°C to be processed for total RNA extraction. Total RNA was extracted from tissue using QIAzol Lysis Reagent (Qiagen). Briefly, tissue samples were minced using dissection scissors and homogenized in QIAzol using Tissue Lyser (Qiagen). Tissue homogenate was phase separated using chloroform, and the aqueous RNA layer was collected and processed using a RNeasy Mini kit to eliminate phenolic impurities carried over from QIAzol extraction. During the column-based clean-up process, on-column DNase treatment was performed to eliminate genomic DNA from the samples. RNA samples were transcribed to cDNA and quantified using a custom porcine Qiagen PCR Array Kit.

### Mouse subcutaneous scaffold implantation

All surgical procedures were approved by Vanderbilt University’s Institutional Animal Care and Use Committee. The ventral region of C57BL/6 mice was shaved and prepped with betadine and ethanol, and small incisions were created, one on either side of the midline. The incisions allowed for the creation of subcutaneous (subQ) pockets. EG2 PTK-UR and EG7 PTK-UR scaffolds were trimmed to 6 x 1 mm discs, UV sterilized, and implanted in the subcutaneous space. The subcutaneous pockets were sutured closed to secure the scaffolds in place.

### Scaffold retrieval and cell isolation

Mice were euthanized 7, 14, and 21 post-surgery, and the scaffolds were explanted from the subQ sites. Scaffold samples were finely minced using dissection scissors in 1 mL PBS. The resultant tissue slurry was digested in a solution of Collagenase D (2 mg/mL), DNase1(0.1 g/mL), and Liberase TL (0.7 mg/mL) for 30 mins at 37°C on an orbital shaker. The digested cell solution was filtered through a 100 µm cell strainer to removed undigested tissue, treated with ACK lysis buffer to remove red blood cells, and washed with cold PBS with 2 mM EDTA and 1% FBS. The cell solution was concentrated into a pellet and resuspended in PBS with 2 mM EDTA and 1% FBS at 1x10^^6^ cells/500 µL. 250 µL of the cell solution was transferred into polystyrene tubes for antibody staining.

### Flow cytometry

Isolated single cells were blocked with Fc block (Invitrogen) for 5 mins and stained for myeloid cell surface markers using the following antibody panel: DAPI, CD45 BriliantViolet 510 F4/80 FITC, CD206 PE, CD11c PerCP-Cy5.5, CD86 PE-Cy7, CD301b eFluor660, and CD11b APC-Cy7 for 30 mins at 4°C. Following staining, cells were washed three times with PBS/1% FBS before being analyzed. All cell populations were gated on live immune cells (DAPI/CD45) prior to phenotype specific gating.

For the lymphoid panel, cells were stained with Live/DEAD Fixable (Invitrogen) followed by a cocktail of surface markers including CD45 BriliantViolet 510, CD8a FITC, TCRγδ PE-Cy7, CD4 APC, and TCRβ APC-Cy7 for 30 min at 4°C. After staining for surface markers, cells were fixed with True Nuclear Transcription Factor Staining Kit for 60 mins at room temperature and stained for FOXP3 PE and GATA3 PerCP-Cy5.5 intracellular markers for additional 30 minutes. Cells were washed three times with PBS/1% FBS before being analyzed. All cell populations were gated on live immune cells (Viability dye/CD45) prior to phenotypic specific gating. Gates were set through the use of fluorescence minus one (FMO) and positive controls. Leukocytes were defined as live CD45+ cells; monocytes and neutrophils were defined as CD45+CD11b+F4/80- population; CD45+CD11b+F4/80+ population (Pan macrophage population) was further analyzed for the presence of CD206 surface marker and CD45+CD11b+F4/80+CD206+ population was isolated. Presence of CD301b surface marker within CD45+CD11b+F4/80+CD206+ macrophage population was quantified and presented as mean fluorescence intensity (MFI) adapted from Shook et al (*38*). Within the CD45+CD11b+F4/80+ macrophage population, we further analyzed the expression of CD11c surface markers and isolated two population, CD45+CD11b+CD11c+F4/80+ (CD11c+ macrophage) and CD45+CD11b+CD11c-F4/80+ (CD11c- macrophage population); dendritic (DCs) cells were characterized as CD45+CD11c+F4/80-. Leukocytes were separated into TCRβ+TCRγδ- and TCRβ- TCRγδ+ to define two lymphoid populations. TCRβ+TCRγδ- were further divided into CD8a+CD4- and CD8a-CD4+ populations to identify cytotoxic T lymphocytes (CTLs) and helper T (TH) cells respectively. The CD8a-CD4+ T helper cell population was used to identify FoxP3+ and Gata3+ regulatory T cells and type 2 helper T cells (TH2). All analyses were performed in FlowJo Flowcytometry Analysis Software (Treetar)

### Analysis

All analyses of qRT PCR data were performed using dCt values calculated from the geometric mean of three housekeeping genes, B2M, HPRT1, and RPL13A. PTK treatment fold change (FC) and log2(FC) were calculated using 2^^-(dCt)^ relative to 900t PE for each time point and presented as geometric mean and arithmetic mean respectively. Heatmaps are presented as row normalized dCt values or log2(FC) as indicated on each plot. For day 31 gene expression comparison, log2(FC) of EG7 PTK-UR is given relative NovoSorb. One-way analyses of variance (ANOVA) followed by Tukey’s pairwise comparison was performed using GraphPad Prism to define statistical differences between treatments at a given time point. Kinetic studies were analyzed using two-way analysis of variance (ANOVA) between groups. Linear regression was used to analyze correlations between EG content and functional readouts including contact angle in vitro, tissue infiltration in vivo, and blood vessel density in vivo.

## Results

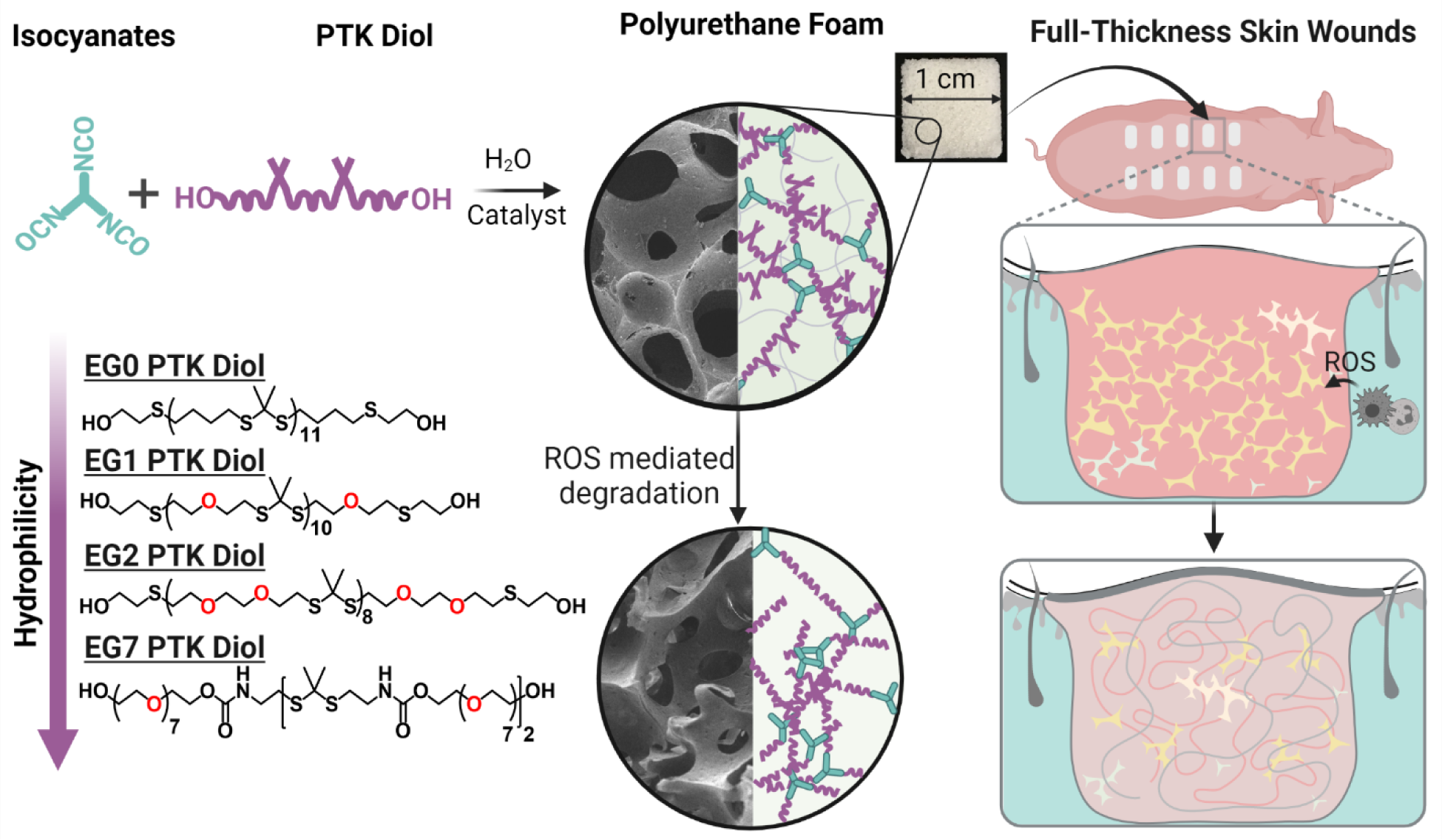

**Abstract:** Reactive liquid molding of isocyanates and poly(thioketal) diols yields covalently crosslinked 3D polyurethane networks with interconnected pores that permit cell infiltration. Structural composition and physical properties of PTK-UR scaffolds can be modulated by controlled variation of the number (0–7) of ethylene glycol units in the PTK diol backbone. PTK-UR foams, which undergo oxidative degradation, act as a provisional matrix for cellular infiltration, vascularization, and ECM deposition for effective repair of skin wounds.

### Synthesis and characterization of PTK diol

EG0, EG1, and EG2 PTK dithiols were successfully synthesized through condensation polymerization of BDT, MEE, and EDDT, respectively, with DMP in the presence of catalytic amounts of pTSA (Figure S1) (*27, 28, 39, 40*). Purified dithiol polymers were effectively functionalized with 2-bromoethanol to obtain EG0, EG1, and EG2 PTK diols. The formation of thioketal bonds was confirmed through ^1^H NMR; δ at 1.6 ppm (Figure S1). Dithiol monomers containing a larger EG content than 2 did not polymerize via condensation polymerization in a controlled manner, likely forming cycles/oligomers (data not shown). To increase the EG content, we developed an alternative polymerization strategy based on polyaddition reaction of heterobifunctional PEG400 nitrophenylcarbonate (HO-PEG-NPC) and a small thioketal diamine crosslinker. The thioketal diamine was synthesized from a protected cysteamine derivative and DMP as confirmed by ^1^H NMR spectroscopy (Figure S2) (*33*). HO-PEG400-NPC was synthesized through the reaction of PEG400 (diol) with p-nitrophenyl chloroformate at a stoichiometric ratio of 1:1 which yielded a mixture of primarily monofunctional PEG400 nitrophenylcarbonate (PEG-NPC) and small percentage of homobifunctional PEG400 dinitrophenylcarbonate (NPC-PEG-NPC) macromers which serve as “chain- extenders” through reaction with multiple thioketal diamines, to obtain EG7 PTK diol polymer (Figure S2). Gel permeation chromatography (GPC) analysis revealed that all PTK diol polymers chain had relatively low dispersity (Ð < 1.2) and number average molecular weight (Mn) of between 1500-2000 gmol^-1^ (Figure S3A). The resulting EG0, EG1, EG2, and EG 7 PTK diols were composed of 11.7, 10.2, 8.2, 1.5 thioketal units per polymer chain respectively (Figure S3C), and the presence of terminal hydroxyl functional groups was verified by their characteristic FTIR absorbance peak at 3400^-1^. Hydroxyl functionalization was quantified indirectly through ^19^F NMR upon reaction with α,α,α-trifluorotoluene. Experimentally determined hydroxyl numbers closely matched theoretical values indicating successful hydroxyl functionalization (>85%) and presence of terminal OH groups for isocyanate reactivity in subsequent foaming reactions (Figure S3C).

The terminal hydroxyl groups on PTK diols were reacted with isocyanate groups present on lysine triisocyanate (LTI) crosslinker in the presence of TEGOAMIN33 as a catalyst and water as a blowing agent to create a polyurethane (PU) foam with an interconnected porous structure; formulation specifications for each foam are provided in Supplementary Table 1. PTK-UR scaffolds were successfully fabricated from EG0, EG1, EG2 and EG7 PTK diols and LTI with similar porosity (86.8% - 91.5%) and comparable pore sizes (74.2-133.3 µm). Efficient covalent crosslinking between OH and NCO-groups of LTI was confirmed based on the low sol fractions for each scaffold (Figure S4).

Contact angle measurements on PU films showed that the EG7 PTK had the lowest contact angle (52.7 ± 5.8°), compared to EG0 (79.2 ± 3.3°), EG1 (66.5± 2.9°), and EG2 (61.2 ± 2.3°) PTK-UR materials describing their relative hydrophilicities (Figure 1A). The measured contact angles positively correlated with the theoretical partition coefficients (Log P) of PTK diols (Figure 1B) and the measured swelling ratios for all of the materials (Figure S4). However, mechanical properties (DMA compressive modulus determined under hydrated conditions) of the scaffolds where all similar with EG0, EG1, EG2, and EG7 PTK-UR scaffolds showing moduli of 75. ± 7.8, 61.4 ± 6.6, 78.5 ± 12.3, and 67.1 ± 10.7 kPa, respectively (Figure 1C); these properties are a good approximation of the reported tensile modulus of skin (*41, 42*). While the wet modulus of all PTK-UR formulations was not significantly different from each other, measurements of the modulus of the 900t and 1500t PE-UR (1500t vs EG0, 1500t vs EG2, p<0.01) scaffolds showed a small but significant decrease compared to the PTK-URs. These collective data indicate that we have generated a series of PTK-UR scaffolds with similar mechanical and structural properties, allowing for careful study of the biological effects of relative scaffold hydrophilicity.

**Figure 1:**
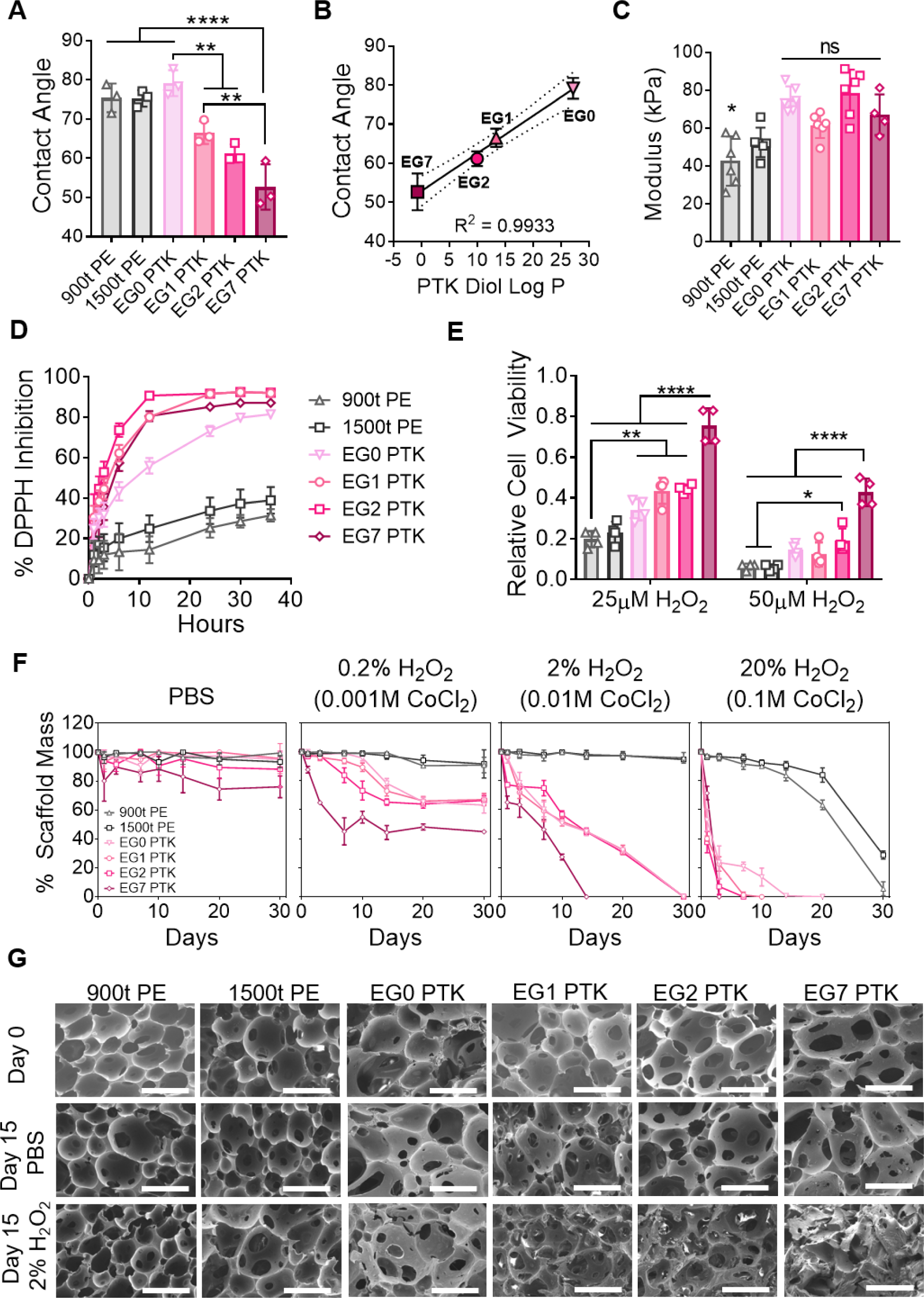
Increasing EG content within PTK diols increases UR scaffold hydrophilicity and radical reactivity. (A) UR film contact angle measurements and (B) correlation between measured contact angles and computed PTK diol Log P values (ChemAxon). (C) UR scaffold mechanical properties measured under compression in hydrated, aqueous conditions. (D) Radical scavenging capacity of PTK-UR compared to PE-UR scaffolds measured through DPPH inhibition. (E) Cytoprotective properties of PTK diols (matched thioketal content) determined after cell exposure to 25 µM or 50 µM of H_2_O_2_ for 24 hours. (F) Scaffold mass loss over 30 day in PBS, 0.2% H_2_O_2_ in 0.001 M CoCl_2_, 2% H_2_O_2_ in 0.01M CoCl_2_, or 20% H_2_O_2_ in 0.1M CoCl_2_ at 37°C. (G) SEM images of PTK-UR scaffolds incubated in PBS or 2% H_2_O_2_/0.01M CoCl_2_ at Day 0 and Day 15; scale bar = 200µm. Data is presented as Mean ± SD, n=3-4 scaffolds, *p<0.05, **p<0.01, ***p<0.001, ****p<.0.0001 by analysis of variance (ANOVA).

### PTK-UR scaffold hydrophilicities positively correlate with EG content and Log P values

TK bonds are broken through irreversible reaction with ROS such as •OH, O_2_ ^-^, H O and ONOO- (*27, 40, 43*). This provides an opportunity to create TK-based biomaterials with inherent antioxidant function and ROS-dependent degradation mechanisms. Using 1,1-diphenyl-2-picrylhydrazyl (DPPH) as a model free radical, we demonstrated the scavenging activity of PTK-UR scaffolds was significantly higher compared to PE-UR scaffolds. The solvent conditions used to perform the DPPH assay (80:20 ethanol/H2O) does not provide an optimal readout for the relative TK bond accessibility in aqueous environments, Despite these conditions, EG1, EG2, and EG7 PTK-UR formulations showed significantly higher radical scavenging compared to EG0 PTK-URs in this polar solvent mixture (Figure 1D). To more directly evaluate the effect PTK composition on antioxidant function and cytoprotection in aqueous environments, we exposed NIH 3T3 murine fibroblasts to varying concentrations of H_2_O_2_ (25µM and 50µM) in the presence of the series of PTK diols (normalized to thioketal molar content). 900t and 1500t PE were treated at 0.5mg/mL solutions to match the highest concentration PTK diol treatment. All PTK diol treatments offered significant protection from H_2_O_2_ (25 µM) mediated cell toxicity compared to 900t PE treatments. At higher concentration of H_2_O_2_ (50 µM), EG7 PTK formulation rescued cell viability to a significantly higher degree with respect to all other treatment groups offering nearly 2-fold greater viability than the next best polythioketal (EG2) (Figure 1E). These data support the ROS scavenging activity of PTK diols and that PTK radical reactivity correlates with hydrophilicity.

To investigate ROS-dependent degradation kinetics of PTK-UR and PE-UR scaffolds, we incubated PU samples in PBS or increasing concentrations of oxidative media (Figure 1F, Figure S5 and S6). PTK-UR scaffolds incubated in hydrolytic media remained intact with minimal mass loss over the course of the experiment. However, unlike 900t and 1500t PE-UR scaffolds, all four PTK-UR foam compositions degraded in oxidative media simulated by hydroxyl radicals (produced from the Fenton reaction between H_2_O_2_ and catalytic amounts of CoCl_2_). The PTK-UR scaffold mass loss was ROS concentration-dependent for all compositions. There were no significant differences in rate of degradation between the different PTK-UR scaffold formulations at the highest hydroxyl radical concentrations. However, at lower concentrations, significantly higher mass loss in EG7 PTK-UR scaffolds was recorded compared to EG0, EG1, and EG2 PTK-UR formulations, despite the EG7 diol having the lowest PTK bond backbone density. Modeling analysis showed that the EG7 PTK-UR scaffolds had a higher degradation rate constant than all of the PTK-UR formations at 0.2 and 2% ROS concentrations and that, under mild ROS conditions (0.2%), there was a correlation between degradation rate constant and EG content (Figure S7, S8 and S9). Morphological changes such as loss in porous architecture and increase in pore size was observed in SEM images of PTK-UR scaffolds incubated in oxidative media, while no apparent changes were visible in PBS incubated samples. Together with the DPPH and cytoprotection assays, these data demonstrate that thioketal groups are more accessible for ROS scavenging in the more hydrophilic (higher EG content) scaffolds.

### Porcine excisional wound model to evaluate structure-functional relationship between scaffold chemistry and wound healing

To investigate the tissue response to synthetic PU scaffolds, we implanted PE-UR and PTK-UR scaffolds in 2 cm^2^ full thickness skin wounds on the dorsal region of Yorkshire pigs. 10 days after scaffold implantation, we observed varying levels of scaffold integration within the wound bed (Figure 2A, top row). A clear increase in tissue integration and homogeneity of distribution throughout the wound bed was observed for the EG7 PTK-UR scaffold when compared to all other treatments (Fig 2A, B). Tissue infiltration within the pores of the scaffolds also positively correlated (R^2^ = 0.92) with EG content in the PTK diol used to fabricate the scaffolds (Figure 2C). The highest tissue infiltration was seen in EG7 PTK treatments (77% ± 3%) compared to analogous PTK chemistries (S8A). Vascular density, visualized through immunohistochemistry (IHC) for vonWillebrand factor (vWF), also positively correlated with EG content (Figure 2A, D, Figure S8B) and thus overall scaffold hydrophilicity

**Figure 2:**
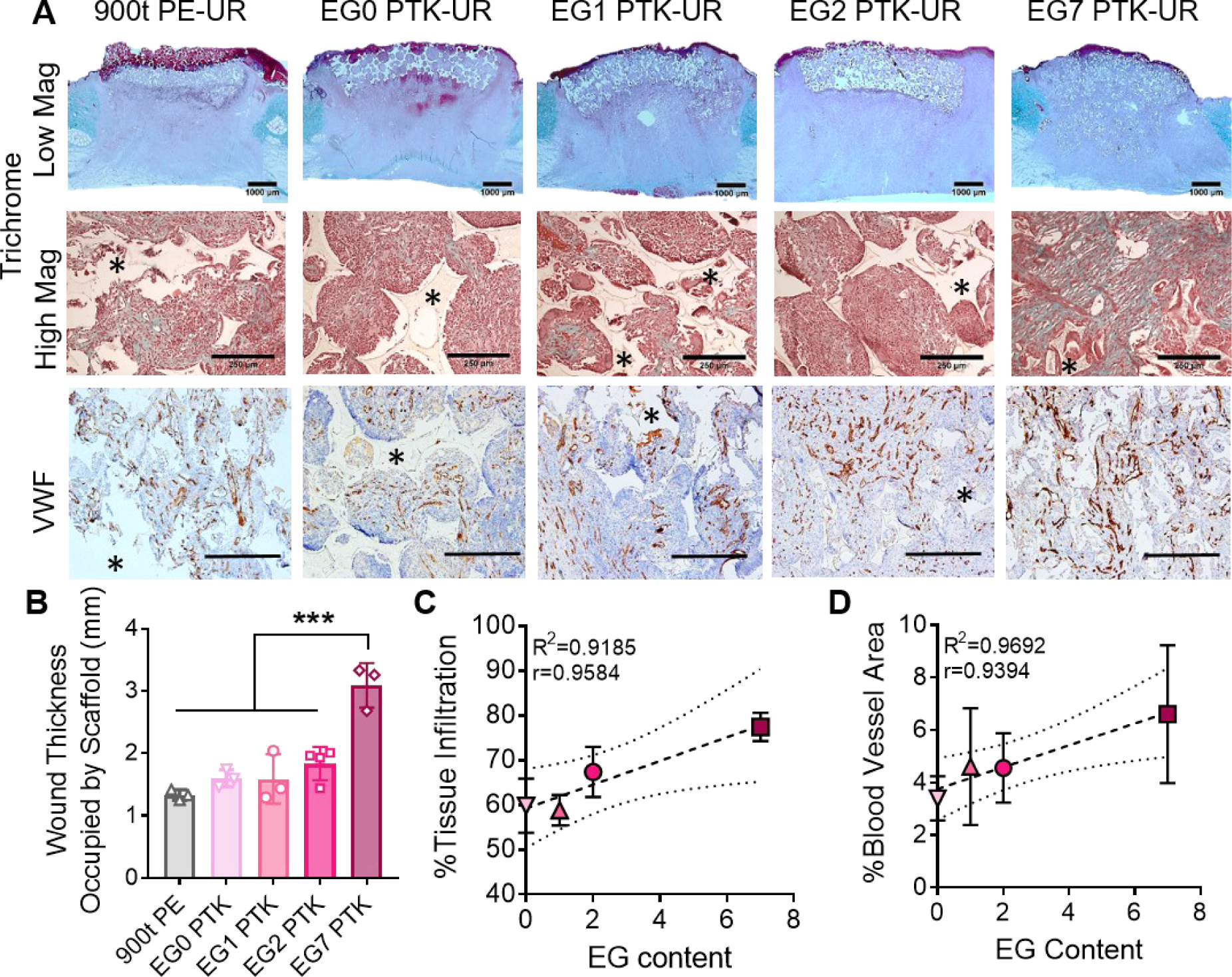
PTK-UR wound bed integration and neovascularization correlate with scaffold hydrophilicity in pig excisional wounds 10 days post implantation. (A) Representative trichrome images showing bulk integration of scaffold within the wound bed (low magnification, scale bar=1000 µm) and cellular infiltration and ECM deposition within scaffolds pores (high magnification, scalebar=250 µm). vWF IHC shows vascularization (brown) within the scaffold-infiltrating granulation tissue (scalebar= 250 μm). (B) Quantification of bulk scaffold integration measured as wound thickness occupied by scaffolds. (C) Positive correlation between PTK diol EG content and percent tissue infiltration within the scaffold pores and (D) PTK diol EG content and blood vessel area; * indicates scaffold remnants. Data presented as Mean ± SD, n=3-4 wounds, ***P<0.001 by analysis of variance

### Scaffold hydrophilicity affects inflammatory cellular infiltrates and microenvironment

Inflammatory response to implanted biomaterials can dictate quality of tissue repair in skin wounds (*44–46*). We investigated spatial distribution of inflammatory cell infiltrates within implanted PU scaffolds 10 days post implantation. All 5 PU scaffolds treatments recruited inflammatory cells including myeloperoxidase (MPO) expressing neutrophils and CD206 expressing macrophages within the scaffolds (Figure 3A). MPO staining was localized around scaffold remnants suggesting recruitment of neutrophils to the scaffold-tissue interface. EG7 PTK scaffolds were associated with significantly fewer neutrophils (Figure 3B) compared to all other treatments. Active recruitment of neutrophils from the blood stream to implanted biomaterials can propagate inflammation through secretion of pro-inflammatory cytokines and interleukins (*47, 48*). Macrophage polarization can also play an important role in biomaterial integration and wound healing (*44, 49, 50*). We investigated the recruitment of M2 macrophages within implanted PU foams and saw a significant increase in CD206 expressing macrophages in EG7 PTK treated wounds compared to 900t PE treatment (Figure 3A, C). To complement histological analysis of wound tissue, we performed bulk tissue analysis of expression of inflammation-associated genes; EG7 PTK-UR scaffold-treated wounds were associated with consistently decreased expression of inflammation related genes including CSF2, CSF3, TNF, IL1B, IL12A, CXCL8, and NOS2 compared to all other treatment groups (Figure 3D).

**Figure 3:**
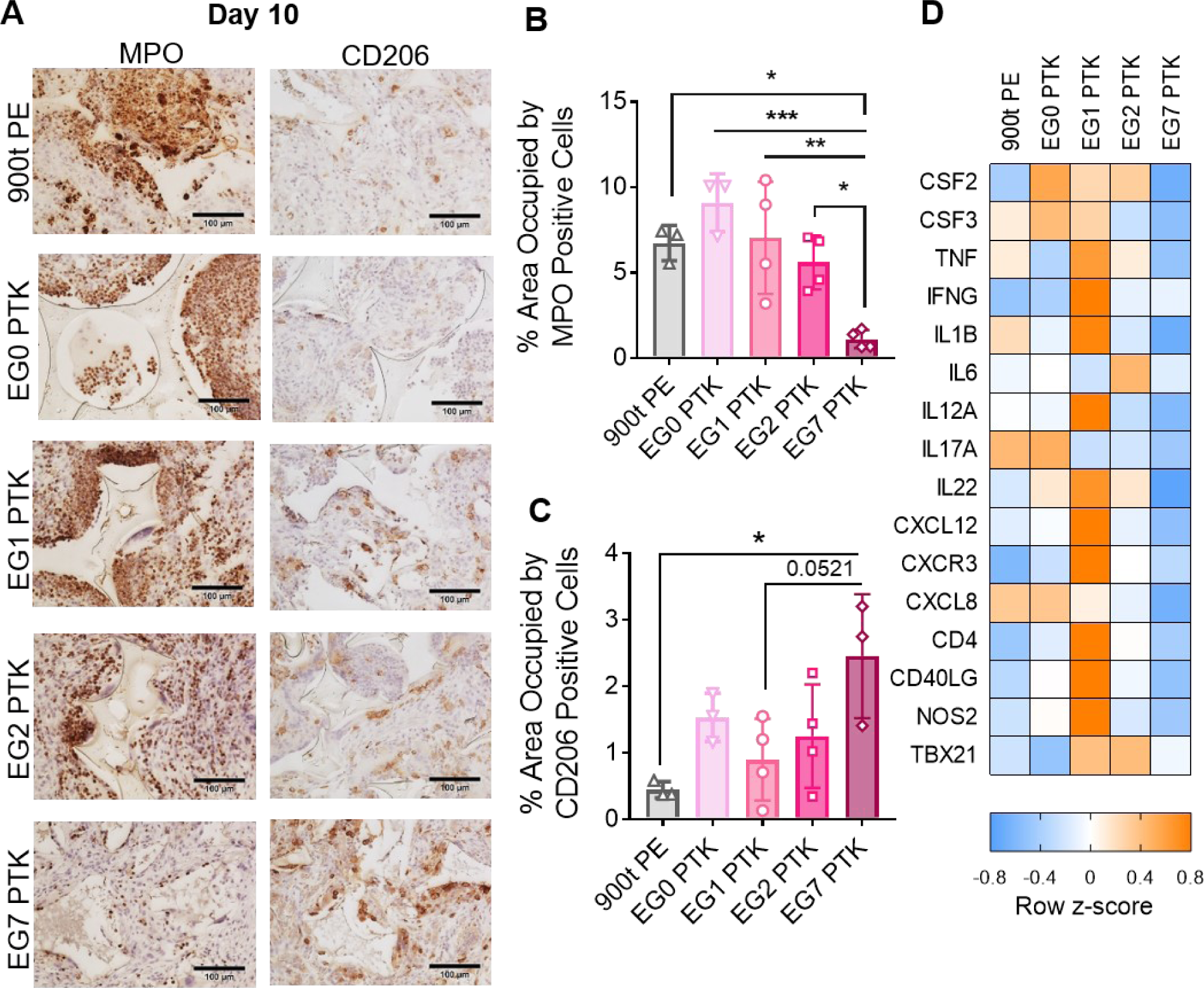
EG7 PTK-UR treated pig excisional wounds have a less pro-inflammatory microenvironment at day 10 relative to more hydrophobic scaffolds. **(A)** Representative MPO (neutrophils) and CD206 (M2 macrophages) IHC images of scaffold infiltrating tissue. Quantified staining at day 10 for **(B)** MPO expressing neutrophils and **(C)** CD206 expressing macrophages. **(D)** Bulk tissue gene expression analysis of inflammatory mediators in porcine skin wound bed 10 days post scaffold implantation. Color-coded heat map showing row normalized z-scores (n=4 wounds per group) of inflammatory marker gene expression. Data presented as Mean ± SD, n = 3-4 wounds, ANOVA *p<0.05, **p<0.01, ***p<0.001. Scale bar = 100µm.

### EG7 PTK-UR scaffolds promote wound closure, re-epithelialization, and overall tissue repair

Next, we investigated wound closure rates of PU foam treated pig skin wounds over a longer time course. Wound area for all scaffold treatments decreased over a period of 24 days (Figure 4A, Supplementary Figure S9A). At 24 days post-surgery, EG7 PTK treated wounds achieved 70.5% closure compared to only 42%, 52.8%, and 53.5% closure observed in EG0, EG1, and EG2 PTK treatments, as well as 49% for 900t PE (Figure 4C). All EG0 PTK-UR scaffolds were excluded from the wounds 20-22 days post implantation resulting in spontaneous wound contraction (Figure S9A). We observed poor EG0 scaffold integration with skin wound despite similar levels of tissue infiltration between EG0 and EG1 scaffolds (Figure S8A and S9B) suggesting the importance of scaffold hydrophilicity in biomaterial-tissue integration. Due to the premature loss of EG0 scaffolds due to extrusion, this treatment group was not included in subsequent analyses.

**Figure 4:**
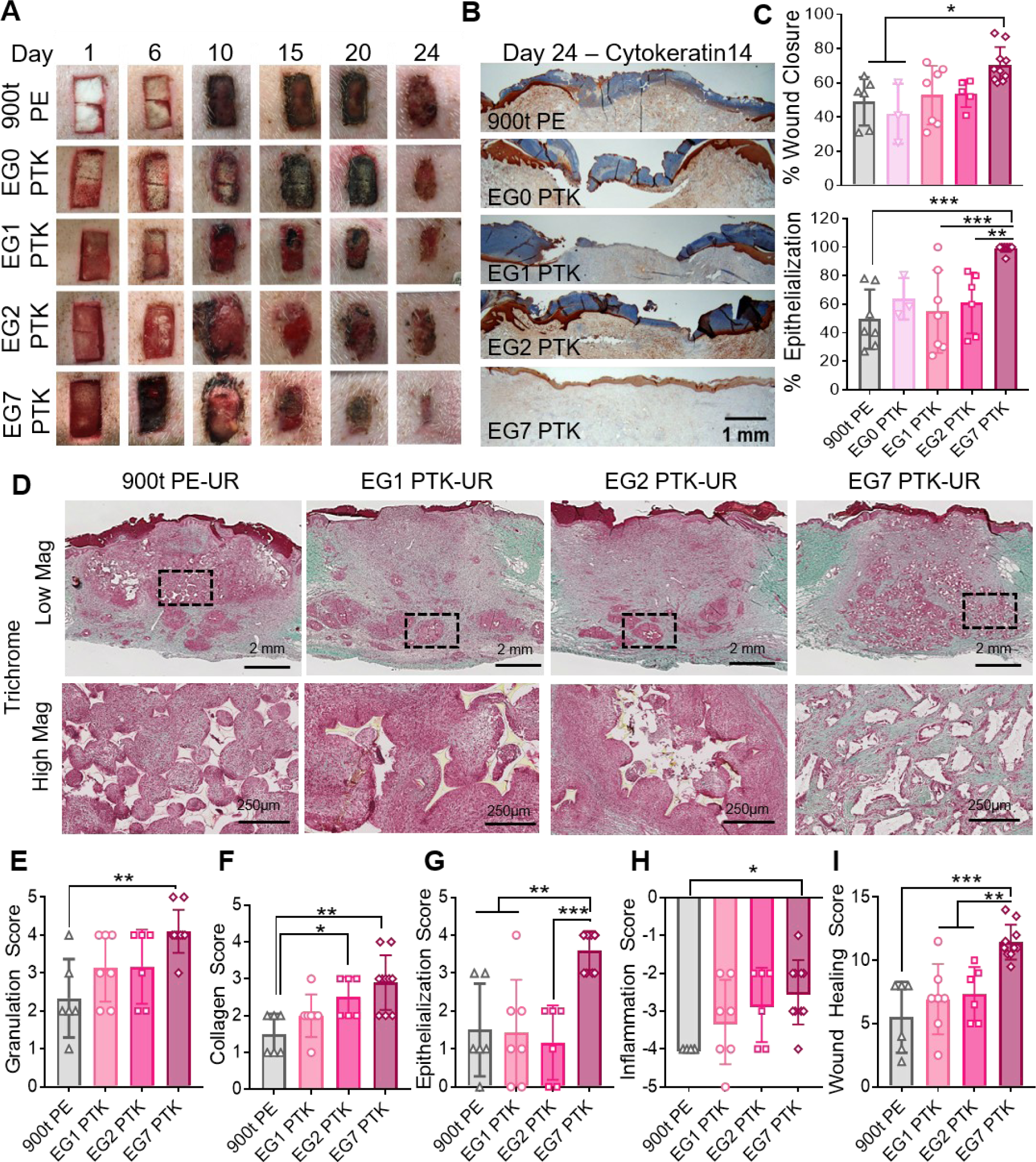
EG7 PTK-UR shows complete re-epithelization and overall enhanced wound healing than more hydrophobic scaffolds in day 24 pig wounds. **(A)** Representative images showing scaffold implantation and wound resolution over a period of 24 days. **(B)** Representative cytokeratin14 IHC showing wound re-epithelialization. **(C)** Wound closure measurements. *Top:* Quantified decreased in wound area (day 24) relative to initial wound size (day 0). *Bottom:* Percentage of wound length covered by a cytokeratin14 positive epidermal layer. **(D)** Representative trichrome images of granulation tissue analyzed 24 days post scaffold implantation (high mag, scale bar= 2mm and low mag, scale bar= 250µm). Wound score was assessed through treatment blinded histopathologist analysis of trichrome and H&E images for **(E)** granulation tissue, **(F)** collagen deposition, **(G)** epithelialization, and **(H)** inflammation, yielding **(I)** a cumulative wound healing score for each scaffold type tested. Data presented as mean ± SD, n = 6-10 wounds, *P<0.05, **P<0.01, ***P<0.001, by analysis of variance (ANOVA)

Complete wound closure is associated with migration of epidermal cells from the surrounding wound margins towards the center, fully covering the granulation tissue with new epithelium. We, therefore, sought to evaluate the re-establishment of epidermal layer in PU treated excisional pig wounds (*51*). Cytokeratin14 IHC, revealed that 90% of EG7 PTK-UR treated wounds had completely epithelialized with 99.2% of the wound covered with a thin, uniform neo-epidermal layer (Figure 4B). On the contrary, none of the other PU treatments resulted in complete re-epithelization of the wounds with an average of 49.6%, 63.9%, 55%, and 61% wound closure observed in 900t PE, EG0, EG1, and EG2 PTK treatments, respectively (Figure 4C). In addition to incomplete epithelization of wounds with these treatments, excessive epidermal thickening suggests persistent inflammation and presence of hyperproliferative keratinocytes with minimal migration (*7, 52, 53*).

Histological analysis of the skin wounds at the day 24 endpoint revealed significant tissue-integration and scaffold resorption across all 4 PU treatments (Figure 4D, low mag.). EG1 PTK and EG2 PTK scaffold remnants were primarily present in the lower portion of the granulation tissue surrounded by a dense layer of immune cells. On the contrary, EG7 PTK-UR scaffolds were uniformly distributed with the granulation tissue and were not associated with dense pockets of immune cells (Figure 4D, high mag). To further investigated the effects of scaffold chemistry on different aspects of skin repair and restoration, trichrome and H&E sections were semi-quantitatively assessed by a treatment-blinded histopathologist. Scoring of the wounds indicated that EG7 PTK-UR treated wounds had higher quality granulation tissue, extensive ECM deposition, and reconstruction of dermal architecture (Figure 4E), along with high quantity of collagen deposition (Figure 4F) compared to wounds treated with 900t PE; there was also a general correlation between EG content and granulation and collagen scores. We also observed significantly higher epithelialization of EG7 PTK treated wounds compared to all other PU treatments (Figure 4G). EG7 PTK treated wounds were associated with lower inflammatory infiltrate, including mononuclear cells and foreign body giant cells (FBGCs) compared to 900t PE, and there was a general inverse correlation between EG content and inflammation score (Figure 4H). The cumulative wound score revealed significantly improved quality of wound healing and repair of EG7 PTK (11.5 ± 1.34) treated wounds compared to more hydrophobic PE (5.5 ± 2.5), EG1 (6.9 ± 2.7), and EG2 (7.33 ± 2.1) PTK formulations (Figure 4I).

### Modulation of innate and adaptive immune response by EG7 PTK-UR scaffolds in pig skin wounds

Both innate and adaptive immune responses can impact biomaterial-tissue integration and wound repair (*48, 54–56*). We hypothesized that increased hydrophilicity of PU scaffolds contributes to decreased pro-inflammatory microenvironment within the wound, ultimately decreasing biomaterial associated fibrosis. We evaluated the effects of scaffold chemistry on the recruitment of immune cells at the scaffold-tissue interface. 24 days post PU scaffold implantation, we confirmed that presence of CCR7+ M1 macrophages and CD3+ T cells indicating the presence of innate and adaptive immune cells (Figure 5A). CCR7 IHC revealed a dense association of M1 macrophages with scaffold remnants, where CCR7 staining intensity decreased as a function of distance from the scaffold-; there was also significantly lower staining surround EG7 PTK scaffold remnants compared to 900t PE, EG1, and EG2 PTK scaffolds at distances greater than 50 µm from scaffold edge (Figure 5B). Unlike EG7 PTK scaffolds, the other treatment groups were associated with a dense layer of M1 polarized macrophages and FBGCs surrounding scaffold remnants within the wound bed, suggesting an ongoing inflammatory response to the scaffold and/or degradation products. To evaluate the adaptive immune response, we also quantified the density of CD3+ T cells surrounding scaffold pieces at day 24. Significantly lower density of CD3 positive T cells was observed surrounding EG7 PTK scaffold remnants compared to EG1 and EG2 PTK scaffolds *in vivo* (Figure 5C). T cells are known to produce IL6, IL12, and IL17 proinflammatory molecules in response to synthetic biomaterial implants, propagating inflammation and fibrosis (*57, 58*). In addition to immune cells, we also looked at αSMA expression in myofibroblasts surrounding scaffolds. We observed structured organization of αSMA expressing myofibroblasts surrounding EG1 PTK and EG2 PTK scaffold fragments but not EG7 PTK scaffolds (Figure 5A). The concentric, spatial organization of M1 macrophages, myofibroblasts, and T cells suggests an active foreign body response against the more hydrophobic PTK-UR scaffold chemistries, similar to that seen in non-degradable implants (*45, 59*). These results suggest that the higher ROS degradability and scavenging potential of the more hydrophilic PTK-UR scaffolds is a critical factor in reducing the immune and foreign body response to the biomaterials. It is also possible that the types of proteins that adsorb preferentially to hydrophilic over hydrophobic material surfaces plays an important role in downstream macrophage polarization (*60–62*) and ultimate healing response.

**Figure 5:**
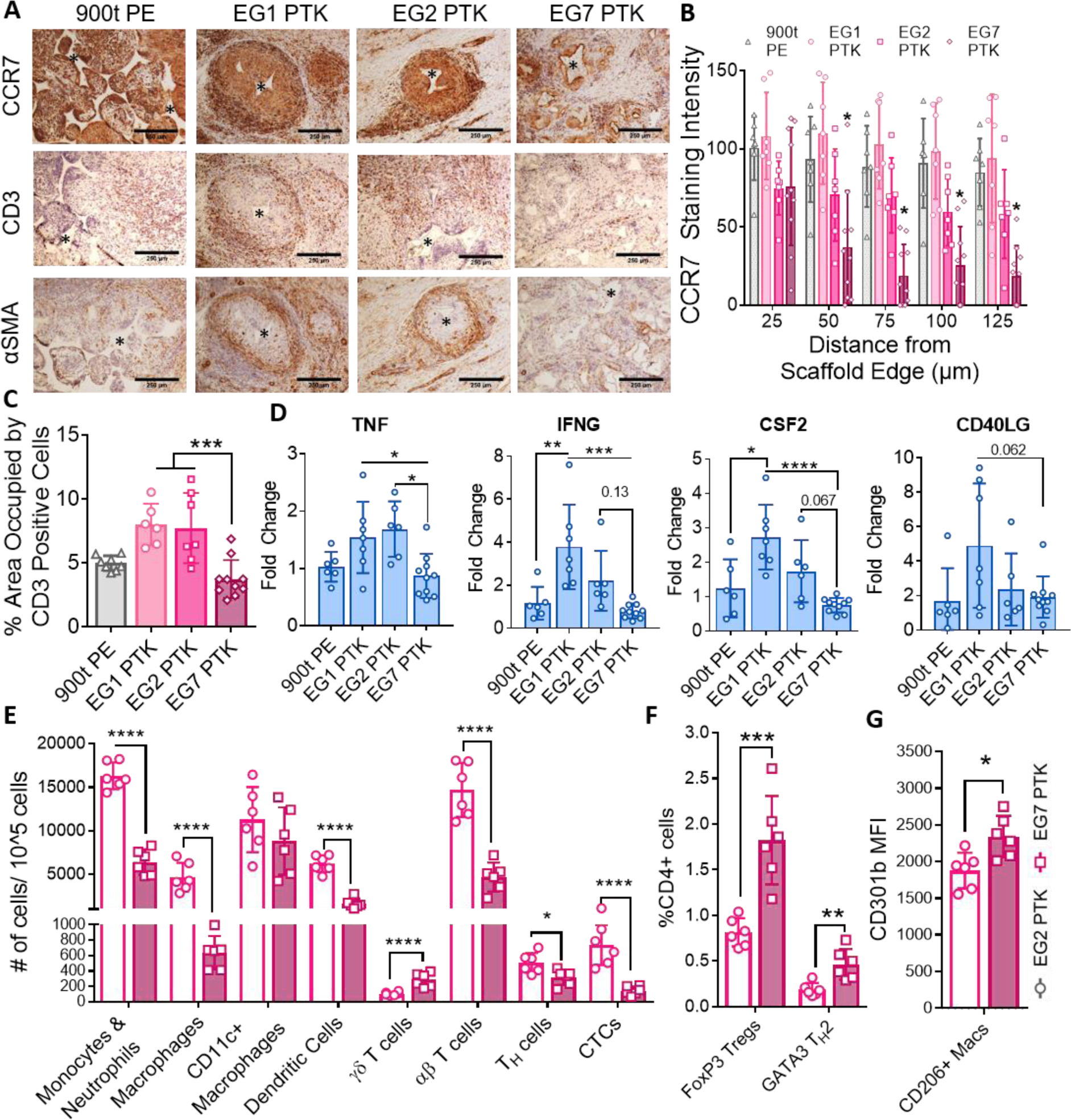
EG7 PTK-UR scaffolds elicit a less pro-inflammatory immune phenotype relative to more hydrophobic variants in day 24 pig skin wounds and day 21 mouse subcutaneous implants. **(A)** CCR7, CD3, and αSMA IHC of pig excisional skin wounds 24 days post treatment. Scale bar = 100 µm. **(B)** Quantification of CCR7 staining intensity as a function of distance from scaffold remnant edges. **(C)** Quantification of CD3 positive pixel area within the wound. **(D)** qRT-PCR quantification of expression of pro-inflammatory genes TNF, IFNG, CSF2, and CD40LG within pig wound scaffolds at day 24. **(E)** Scaffold infiltrating myeloid and lymphoid populations quantified 21 days post subcutaneous implantation of EG2 PTK (empty bars) and EG7 PTK (pink filled bars) scaffolds in mice. **(F)** Percentage of FOXP3+ and GATA3+ infiltrating CD4 helper T cells in EG2 vs EG7 scaffolds. **(G)** Expression of CD301b in EG7 scaffold infiltrating CD206+ macrophages compared to EG2 PTK scaffolds. Data presented as Mean ± SD, n=6-10 wounds, *P<0.05, **P<0.005, ***P<0.001, ****P<0.0001 by analysis of variance (ANOVA)

To further evaluate the inflammatory microenvironment within PU scaffold treated wounds, we quantified bulk tissue expression levels of proinflammatory genes (Figure 5D). We observed significantly lower expression of tumor necrosis factor-α (TNFA), a pro-inflammatory cytokine, in EG7 PTK treated wounds compared to EG1 and EG2 PTK treatments. Similarly lower levels of expression were seen for proinflammatory mediators, interferon-γ (IFNG) and colony-stimulating factor 2 (CSF2). CD40LG, a costimulatory-marker expressed on activated T-cells also trended toward being lower in EG7 PTK treated wounds compared to EG1 PTK (p=0.062).

Due to loss in spatial resolution of inflammatory regions in bulk tissue expression analysis and limited availability of porcine specific antibodies for IHC, we utilized a mouse subcutaneous model to further characterize the immune cell populations and phenotypes that infiltrate the scaffolds. This flow cytometry study (gating strategies outlined in Supplement Figure 10) focused on EG2 and EG7 PTK-UR scaffolds, as these materials both effectively integrate in vivo while still showing significant hydrophilicity-related, biological differences. Scaffolds were implanted into ventral subcutaneous pockets in C57BL6 mice, and scaffold infiltrating cells were analyzed after 1, 2, and 3-weeks post implantation. After 1-week, we observed similar levels of CD45+ leukocytes, CD11b+CD11c-F4/80+ macrophages (20% of all leukocytes), CD11b+F4/80- monocytes/neutrophils (40% of all leukocytes), and CD11c+F4/80- dendritic cells (15% of all leukocytes) within the EG2 and EG7 scaffolds (Fig S11). Over a period of 3 weeks, infiltrating leukocyte populations significantly decreased in EG7 PTK scaffolds but remained constant in EG2 PTK-UR scaffolds, suggesting an unresolved inflammatory response. We observed a significant decrease in CD11b+CD11c-F4/80+ macrophage population in EG7 PTK scaffolds compared to EG2 scaffolds. Similar decreases in CD11b+F480- monocyte and neutrophil populations were also observed. After 3 weeks of scaffold implantation, we also observed significantly lower number of myeloid populations including CD11b+F480- monocyte and neutrophil, CD11b+CD11c-F4/80+ macrophages, CD11c+F4/80- dendritic cells (DCs), αβ+ T cells, CD4+ helper T cells, and CD8+ cytotoxic T lymphocytes (CTLs) present in EG7 PTK cellular infiltrate compared to EG2 PTK (Figure 5 E). On further analysis of T cell phenotypes, we saw a significantly higher percentage of CD4+FoxP3+ regulatory T-cells (T-regs) and CD4+GATA3+ type-2 helper T cells (TH2) within EG7 PTK scaffolds compared to EG2 PTK-UR scaffolds (Figure 5F). We also detected a higher number of γσ T cells associated with EG7 PTK scaffolds; these cells are a source of fibroblast growth factor 9 (FGF9) and keratinocyte growth factor (KGF), which are both known to support re-epithelialization and hair follicle neogenesis after wounding (*63, 64*). We observed significantly higher expression of CD301b on CD206+ macrophages isolated from EG7 PTK compared to EG2 PTK scaffolds (Figure 5G). CD301b+ macrophages support effective skin wound healing, and CD301b is used as a marker for isolation of pro-regenerative macrophages from injured tissue (*65–67*).

### EG7 PTK-UR dermal substitute enables effective tissue repair

We next measured effects of scaffold chemistry on wound expression of genes related to granulation tissue formation, proliferation, and remodeling processes of wound healing in day 24 pig wounds (Figure 6). We observed that EG7 PTK scaffold treated wounds had approximately 4-fold higher expression of genes encoding ECM proteins that comprise granulation tissue, including collagen I, III, V, and XIV (COL1A2, COL3A1, COL5A2, COL14A1) and tenascin (TNC). Integrins such as integrin β1 (ITGB1), integrin β3 (ITGB3), and integrin αv (ITGAV), known to play important roles in cell adhesion, migration, and proliferation (*68, 69*) were also significantly up regulated in EG7 PTK treatments. In addition, matrix metalloproteinases (MMP) including gelatinases (MMP2 and MMP9) and collagenases (MMP7), known to play important roles in collagen remodeling and recruitment of endothelial cells for neovascularization (*70, 71*) were significantly upregulated in response to EG7 PTK treatments with respect to other PE and PTK treatments. Significantly lower expression of MMP1, a type 1 collagenase upregulated in chronic wounds (*72*), was observed to be more lowly expressed in EG7 samples relative to EG1 and EG2 PTK treatments. We also observed an increase in TIMP2, an important regulator of MMPs produced by basal keratinocytes, in EG7 PTK treated wounds, suggesting that a homeostatic balance between MMPs and their inhibitors (TIMPs, tissue inhibitor of metalloproteinases), existed in EG7 treated wounds (*73*). Transforming growth factor β (TGF-β) pathway related genes such as SMAD3 and TGFB3, which are known to promote collagen biogenesis and polarization of macrophages to M2 pro-healing phenotype, were significantly upregulated in EG7 PTK treated wounds compared to other treatments (*74*). Similar trends of increased gene expression were observed in EG7 samples for genes such as insulin like growth factor (IGF), fibroblast growth factor 7 (FGF7), connective tissue growth factor (CTGF), and platelet derived growth factor (PDGF), all factors that promote cell proliferation and migration to drive skin wound granulation, vascularization, and re-epithelialization (*75*).

**Figure 6:**
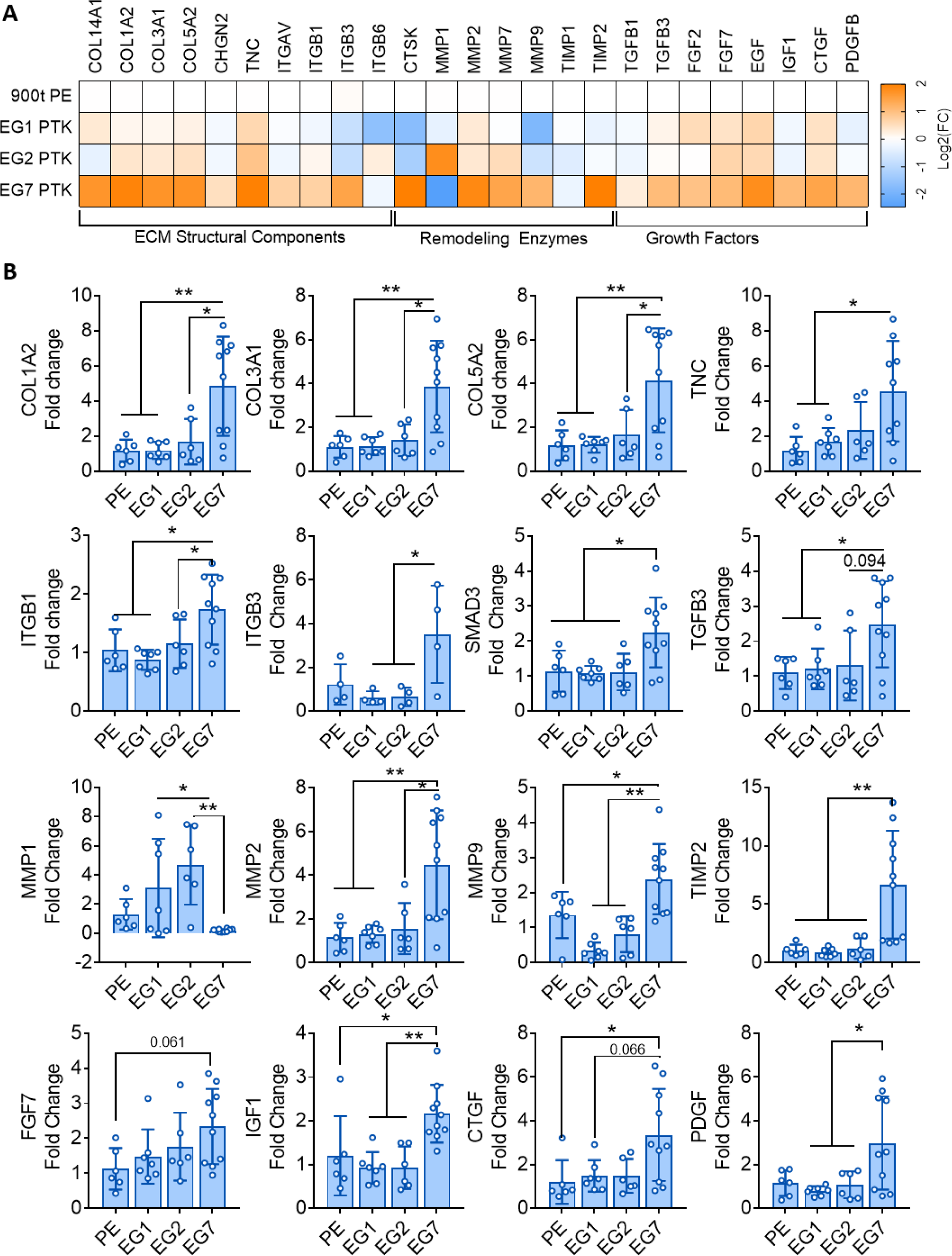
EG7 PTK-UR scaffolds promote higher expression of pro-growth, ECM, and remodeling genes relative to more hydrophobic scaffolds in day 24 porcine wounds. **(A)** Bulk tissue differential gene expression of genes encoding ECM components, remodeling enzymes, and growth factors 24 days post-wound. Gene expression was analyzed and displayed as color-coded heat map showing log_2_(FC) relative to PE 900t control. **(B)** Expression fold change (FC) of selected genes relative to wound healing. Data presented as Mean ± SD, n= 4-10 wounds, *P<0.05, **P<0.01, **P<0.001, by analysis of variance (ANOVA)

### Benchmarking EG7 PTK-UR against clinically approved dermal substitutes

The EG7 PTK-UR scaffold formulation was tested in pig skin wounds relative to clinically approved dermal substitutes including collagen-based Integra Bilayer Wound Matrix (BWM) and polyester-based polyurethane NovoSorb Biodegradable Temporizing Matrix (BTM). We implanted porcine full thickness skin wounds with either EG7 PTK, NovoSorb, or Integra and measured wound closure over time and tissue histological outcomes at 31 days post implantation. All 3 dermal substitutes facilitated tissue infiltration and wound closure over a period of 31 days (Figure 7A, Supplemental Figure 12A). Unlike Integra, EG7 PTK scaffolds provided optimal wound stenting for the first 10 days of wounding similar to Novosorb. Following initial wound stenting, EG7 PTK treated wounds resolved at a similar rate to Integra with no significant differences in the rate of wound closure. However, we observed significant increases in the rates of wound closure in EG7 PTK and Integra vs NovoSorb at days 14, 17, 19 and 21 post implantations. Upon removal of the non-degradable polyurethane layer (according to manufacturer guidelines), the wound area of NovoSorb treated wounds significantly decreased, suggesting rapid loss of stenting provided by the protective layer, followed by consequent wound contraction (Figure 7B). In addition to wound closure, we measured vascular perfusion of scaffold treated wounds using laser doppler perfusion imaging (Supplemental Figure12B). At earlier time points (day10 and day17), EG7 PTK-UR treated wounds were significantly more perfused than Integra and NovoSorb suggesting higher neovascularization within the wounds (Figure 7C). Removal of temporary epidermal layer constructs from Integra and NovoSorb (Figure 7B, indicated by arrows) resulted in disruption of granulation tissue causing re-wounding and increased perfusion within the wounds at days 17 and 24 respectively.

**Figure 7:**
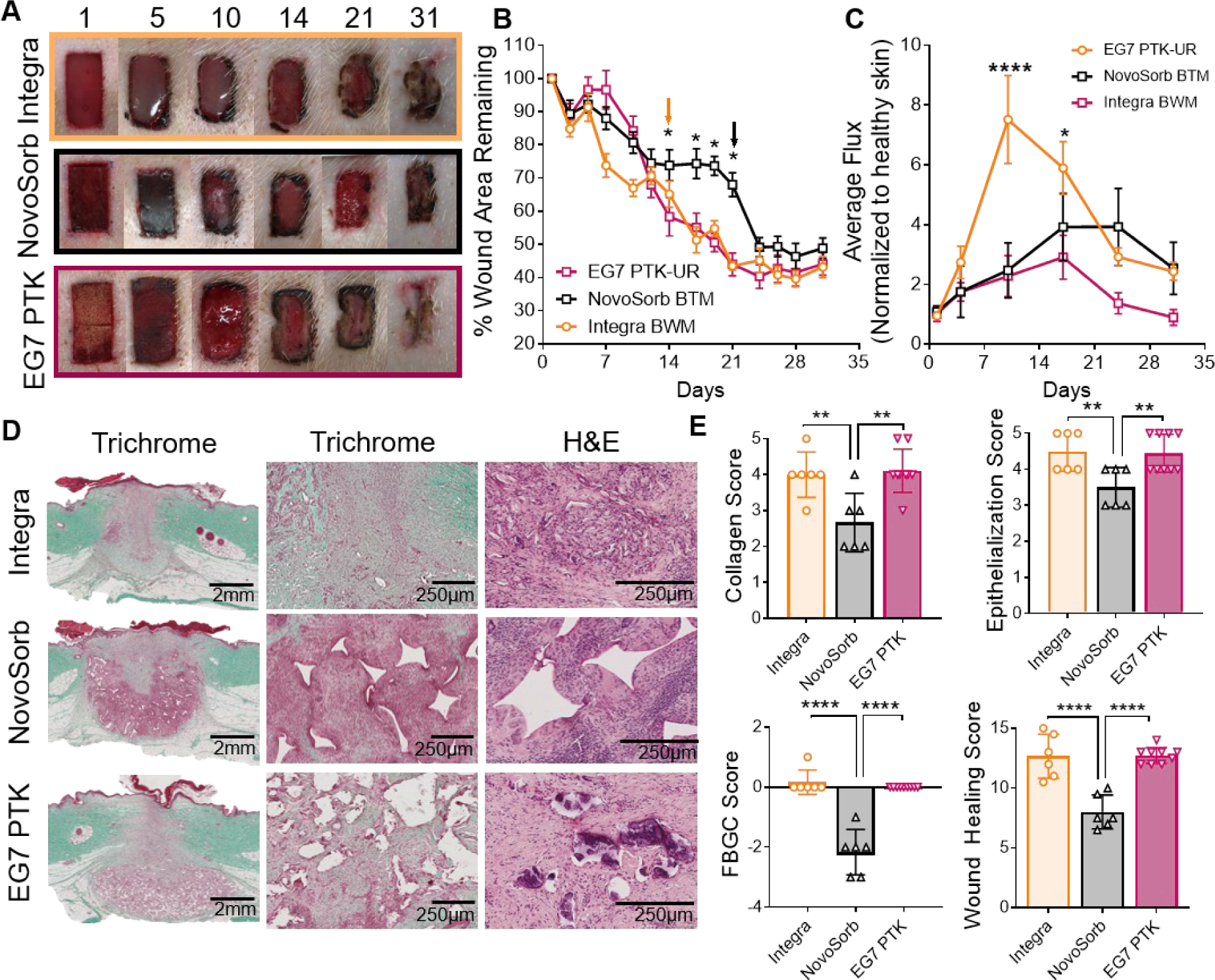
EG7 PTK-UR scaffolds wound healing benchmarking versus Integra BWM and NovoSorb BTM in day 31 pig excisional wounds. **(A)** Images showing scaffold implantation and temporal closure of a 2 x1 cm porcine skin wound treated with collagen-based Integra BWM, polyester based NovoSorb BTM, and thioketal based EG7 PTK-UR. **(B)** Kinetics of wound closure; arrows indicate timepoint of manufacturer-recommended removal of protective layer from Integra and NovoSorb. **(C)** Relative blood perfusion within scaffold-bearing wounds measured by LDPI. **(D)** Trichrome and H&E images of wound sections 31 days post treatment with different dermal substitutes showing quality of infiltrating tissue and residual cell response to scaffold remnants. **(E)** Semi- quantitative analysis of wound healing through treatment-blinded pathohistological scoring of trichrome and H&E images for collagen deposition, epithelialization, and foreign body giant cells (FBGC), yielding a cumulative wound healing score. Data presented as Mean ± SEM (Fig B, C) and Mean ± SD (Fig E), n = 6-10 wounds, *P<0.05, **P<0.01, ***P<0.001, ****P<0.0001, by analysis of variance (ANOVA)

Histological examination of scaffold treated wounds revealed formation of granulation tissue along with the reestablishment of epidermal layer in all 3 treatment groups. Trichrome and H&E stained sections revealed dense cellularity around NovoSorb scaffold remnants along with the formation of FBGCs otherwise minimally present or absent in EG7 PTK treated wounds (Figure 7D). EG7 PTK-UR treated wounds were composed of denser, more organized collagen deposition, along with decreased cellularity, arounds scaffold remnants. This observation indicates that the EG7 PTK-UR treated wounds were more mature and had transitioned into a more advanced remodeling phase relative to Novosorb treated wounds. Treatment-blinded histopathological scoring of these sections revealed similar collagen deposition, re-epithelialization, and FBGC density between Integra and EG7 PTK treated wounds (Figure E). In contrast, NovoSorb treated skin wounds showed significantly lower collagen deposition with loosely arranged fibers, incomplete re-epithelialization, persistence of inflammatory infiltrate, and formation of FBGCs. Cumulative wound healing score determination revealed significantly decreased quality of wound healing of NovoSorb treated wounds compared to Integra and EG7 PTK-UR treatments, with no significant differences detected between the latter two materials.

### Effective dermal repair and immune modulation by EG7 PTK-UR treatments compared to NovoSorb

Next, we more thoroughly characterized the wound microenvironment of EG7 PTK-UR scaffold treated pig wounds versus the more structurally analogous, synthetic Novosorb PU foam treated wounds. To complement the LDPI data, we measured vasculature within scaffold-treated wounds using vWF IHC and observed significantly higher blood vessel density within EG7 PTK treated wounds compared to NovoSorb (Figure 8A, B). To further examine the differences in dermal repair and remodeling between the two skin substitutes, we analyzed expression of genes involved in ECM deposition and remodeling along with growth factors implicated in dermal wound healing. Gene analysis revealed significant upregulation of COL1A2, COL3A1, COL5A2, COL14A1 and TNC in EG7 PTK treated wounds compared to NovoSorb (Figure 8C). Higher levels of TNC, ITGB1 and ITGB6 were also observed. Additionally, significantly higher levels of ECM remodeling enzymes including cathepsin K (CTSK), MMP2, MMP7 and MMP9 and TIMP2 were seen in EG7 PTK treated wounds compared to NovoSorb. Factors involved in collagen biosynthesis and ECM remodeling such as TGFB3 and SMAD3 were also significantly higher in EG7 PTK treated wounds. Genes encoding various growth factors including FGF2, FGF7, epidermal growth factor (EGF), IGF1 (insulin like growth factor 1), CTGF and PDGF implicated in endothelial, fibroblast and, keratinocyte proliferation and migration were significantly higher in EG7 PTK treated wounds compared to NovSorb at day 31 post treatment (*75*).

**Figure 8:**
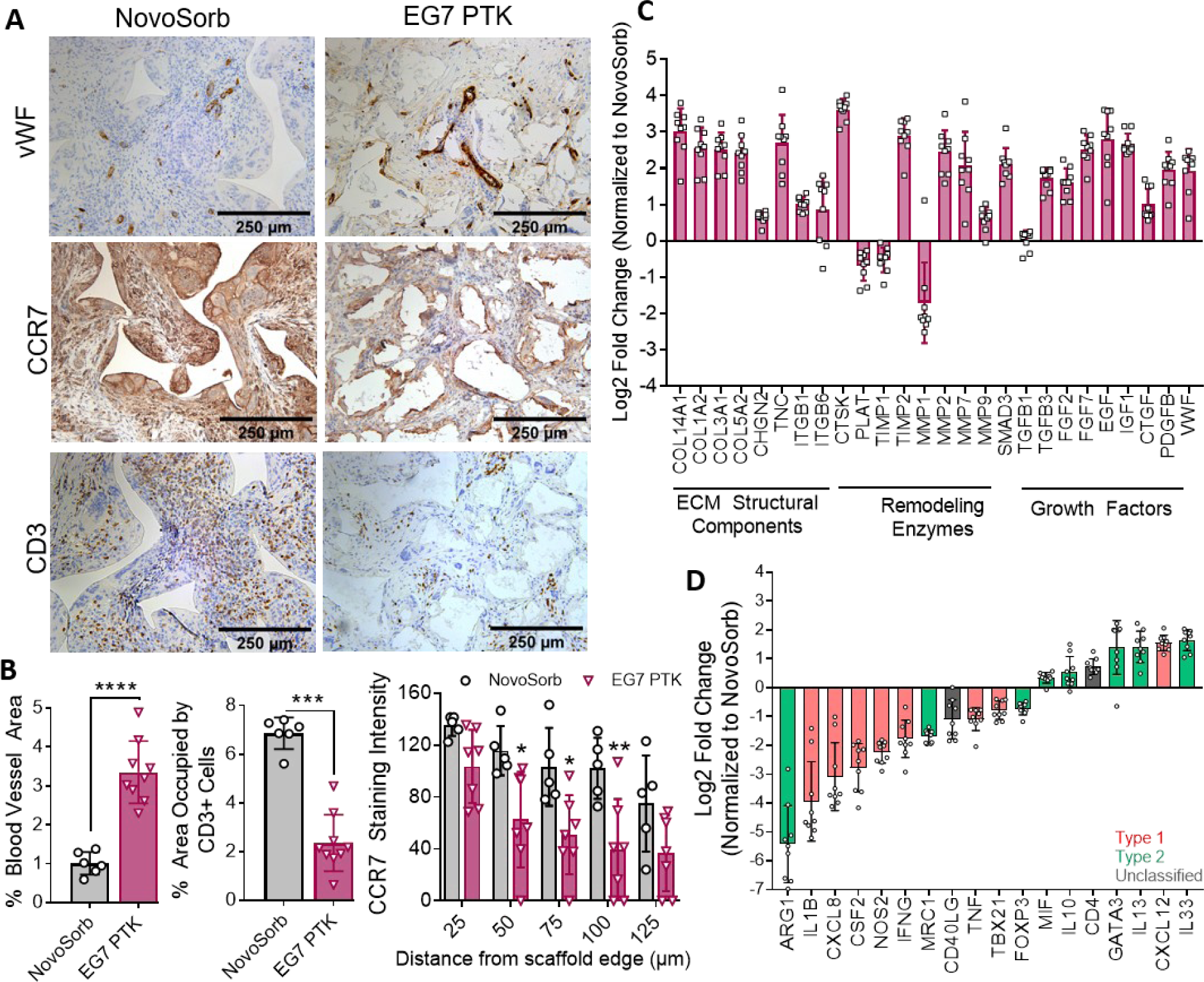
EG7 PTK-UR scaffolds induce more vascularization and higher expression of wound healing-related and anti-inflammatory genes in day 31 pig wounds compared to NovoSorb. **(A)** vWF, CCR7, and CD3 IHC images of NovoSorb and EG7 PTK-UR treated wounds. **(B)** Quantification of vWF, CD3, and CCR7 positive pixels using ImageJ. **(C)** Relative expression of genes encoding ECM components, remodeling enzymes, and growth factors related to wound healing and remodeling. **(D)** EG7 PTK-UR-treated wound expression of pro-inflammatory (red) and anti-inflammatory genes (green) (relative to NovoSorb). Data presented as mean ± SD, n = 6-10 wounds, *P<0.05, **P<0.005, ***P<0.001, by analysis of variance (ANOVA).

Deeper analysis of innate and adaptive immune cells within scaffold infiltrating tissue showed significantly lower density of CD3+ T cells in EG7 PTK treated wounds compared to NovoSorb suggesting resolution of the inflammatory wound healing phase along with decreased inflammation surrounding scaffold remnants and degradation products (Figure 8A, B). Concurrently, we observed significantly lower density of CCR7+ macrophages and FBGCs associated with PTK-UR scaffold remnants compared to NovoSorb, with CCR7 staining intensity varying as a function of distance from scaffold edge (Figure 8A, B). To further characterize the inflammatory response, we performed qRT-PCR on bulk wound tissue samples. Relative to NovoSorb, EG7 PTK-UR samples were associated with significantly lower expression of pro-inflammatory genes such as interleukin 1β (IL1B) and TNF. A similar decrease in expression of inflammatory chemokine ligand 8 (CXCL8) and CSF2 as well as M1 macrophage marker nitric oxide synthase 2 (NOS2) and TH1 specific transcription factor T-bet (TBX21) were observed within EG7 PTK treated compared to NovoSorb. Decreases in pro-inflammatory markers in EG7 PTK treated wounds were accompanied by a concomitant upregulation of anti-inflammatory markers such as TH2 transcription factor GATA3 and interleukins including IL10, IL13, and IL33 that have been implicated in type 2 immunity, tissue regeneration, and wound healing (*76–79*).

## Discussion

Chronic skin wounds are growing in prevalence (affecting nearly 4.5 million Americans) and have a 40% chance of recurrence (*3*), emphasizing the need for low cost and efficacious, degradable dermal substitutes. Naturally-derived biologic scaffolds face issues with reproducibility and costs of production, while synthetic scaffolds, especially PU-based foams, are a promising alternative that is more economic to manufacture. The current study was designed to investigate the impact of foam hydrophilicity on wound healing outcomes and facilitated the identification of a new, leading PTK-UR scaffold formulation (EG7). The EG7 formulation is more hydrophilic than any PTK-UR foams generated previously and showed high potential in promoting repair in clinically-relevant porcine wounds, a model system that provides higher congruency to human skin wounds relative to common rodent models (*80*). The EG7 formulation also outperformed a predicate synthetic scaffold device and performed similarly to a well-established, naturally-derived scaffold with a history of clinical wound healing benefits but high associated product cost (*15*).

Tissue damage and biomaterial implantation are both consistently associated with initial extravasation of circulating neutrophils, monocytes, and macrophages to the site of insult (*81*). These inflammatory first responders generate reactive oxygen species (ROS) such as hydroxyl radical (•OH), hydrogen peroxide (H_2_O_2_), hypochlorite (ClO-), and superoxide (O2-) as part of the innate immune response. In the current work, ROS are harnessed as biological, cell-mediated degradation stimuli for PTK-UR scaffolds (*8*). In turn, the PTK-UR scaffolds have antioxidant function, which can protect the healing tissue from oxidative stress, which is a consistent feature of chronic wounds due to persistent type 1 immune response (*82–84*). Unlike cell-responsive PTK scaffolds, polyester-based urethane scaffolds (PE-UR) undergo non-specific hydrolysis, releasing acidic byproducts that can trigger autocatalytic degradation and rapid, mid- to late-state material loss and consequent pore collapse, inhibition of cell infiltration, and poor wound stenting (3). To overcome the shortcomings of PE-URs and to better elucidate structure-function relationships of PTK- URs, we synthesized a series of PTK diols with varied numbers of EG repeats between TK bonds in the polymer backbone. These series of PTK diols enabled a systematic evaluation of the effect of hydrophilicity on ROS mediated scaffold degradation, immune response, vascularization, and quality of tissue repair.

While assessing the oxidative degradation of PTK-UR scaffolds *in vitro*, it was noted that a decrease in thioketal content between EG0 and EG7 (11.2 vs 2) PTK diols did not result in a proportional decrease in degradation rate (0.6 s^-1^ vs 1.4 s^-1^, EG0 vs EG7) (Supplemental Figure 3C, 7B). Instead, a higher rate of degradation and mass loss was seen in EG7 PTK scaffolds when exposed to 0.2% and 2% H_2_O_2_/CoCl_2_, highlighting the importance of scaffold hydrophilicity on the rate of (oxidative) degradation. This is likely due to increased exposure and susceptibility of thioketal bonds to cleavage by strongly polar oxidative species in a hydrated environment. The differences in radical scavenging potential between PTK-UR chemistries was diminished in organic solvents (DPPH assay performed in 80-20% EtOH: H2O) further highlighting the importance of having a well solvated PTK bond to increase ROS reactivity. It should be noted that, despite a similar overall EG content in EG2 and EG7 PTK scaffolds (20 vs 14 EG repeats per diol), significant differences in swelling ratio (200% vs 600% for EG2 and EG7, respectively) were observed. This is likely due to the inherent hydrophobicity of thioketal bonds and highlights the importance of having strongly hydrophilic groups that serve as spacers between the thioketals along the polymer backbone to ensure accessibility of the TK bond within aqueous environments. Interestingly, computation of theoretical Log P values indicated that EG7 PTK diols have a Log P value below zero (indicating hydrophilicity and a partition preference of water over octanol), whereas the remainder of the EG series were greater than 1, indicating their net hydrophobic character.

Differences in scaffold-tissue integration between EG0/EG1/EG2 PTK-UR scaffolds and EG7 PTK-URs were pronounced when implanted in porcine excisional skin wounds. Despite similarities is porosity, pore size, and dynamic mechanical compressive properties, all physical features that affect tissue response (*44, 85, 86*), EG7 PTK-UR scaffolds were more evenly distributed throughout the thickness of the wound (3.1 ± 0.4 mm) while there was lower tissue integration (< 2mm) of more hydrophobic scaffolds (Figure 2A, B). EG7 PTK-UR scaffolds, likewise, had greater vascularization, which may be attributable to relatively low expression of myeloperoxidase (Figure 3A, B), an antiangiogenic enzyme highly expressed in chronic wounds (*87*), and higher expression of growth factors PDGF (*88, 89*) and TGFB3 (*90*) (Figure 6B), which positively impact endothelial cell proliferation, migration, and vessel maturation. The more desirable wound healing response of the EG7 vs more hydrophobic PTK-UR scaffolds is potentially linked to mechanisms such as improved biological response to the more swollen scaffold architecture, the composition of the protein adsorbate (*61, 91*), better ROS reactivity and scavenging (Figure 1E), and more rapid breakdown and better clearance of more hydrophilic degradation products.

Wound closure by re-epithelialization is a key aspect of wound healing that is typically a primary outcome in clinical studies because re-establishment of skin epidermal barrier is associated with reduced risk for infection and patient morbidity. A moist environment within chronic wounds has been linked with positive healing outcomes, as opposed to slower rates of wound closure and incomplete resurfacing seen in dry wounds (*92*). The ability of the EG7 PTK-UR to maintain a more hydrated wound environment may also contribute to the significantly higher epithelial resurfacing of EG7 PTK treated wounds compared to more hydrophobic foams (Figure 4C). In addition to the physical properties of EG7 PTK-UR foams, these scaffolds also recruited significantly higher (2.7-fold) numbers of γδ T cells compared to EG2 PTK scaffolds when implanted in of the mouse subcutaneous space (Figure 5E). This subpopulation of T cells is present in the epidermal layer where it coordinates epithelial proliferation and migration through secretion of growth factors (*64*). Significant upregulation of IGF1 (2-fold), TGFB3 (2-fold), and PDGF (3-fold) in EG7 PTK-UR scaffold treated wounds was also observed, genes anticipated to promote keratinocyte migration and re-epithelialization (*51*).

Relative PTK-UR scaffold hydrophilicity also had a significant impact on the surface response to scaffold remnants in the later stages of wound healing in terms of the composition and phenotype of macrophages, T cells, and myofibroblasts (Figure 5A), cell types responsible for non-degradable implant fibrous encapsulation (*57, 59, 93*). Active recruitment of neutrophils, monocytes, macrophages, and dendritic cells to the site of biomaterial implantation can propagate inflammation and foreign body response resulting in tissue and implant structural damage (*94*). Hydrophilic and anionic surfaces can cause biomaterial-adherent macrophage apoptosis and lower macrophage fusion, potentially limiting the deleterious effects of pro-inflammatory macrophages and reducing formation of FBGCs (*95*), as supported by our observation that EG7 PTK scaffolds were associated with significantly fewer CCR7+ M1 macrophages and FBGCs relative to more hydrophobic PTK-URs (Figure 5 B). Dendritic cells (DCs) can serve as an important bridge between innate and adaptive immunity by interacting with biomaterials through pathogen recognition receptors (PPR), leading to downstream antigen presentation and activation of T cells. Hydrophilic surfaces can also limit DC adhesion and maturation and consequently reduce pro-inflammatory T cell response (*96, 97*). Indeed, subcutaneous implant mouse studies showed that EG7 PTK scaffolds had decreased number of infiltrating DCs compared to the more hydrophobic EG2 PTK-UR. These collective data suggest that the more hydrophobic scaffolds have a stronger and more sustained immune response compared to EG7 PTK-UR foams and that this facet has implications in biomaterial wound healing potential.

To establish the therapeutic potential of the EG7 PTK-UR scaffolds more broadly in wound healing, we benchmarked against commercially available Integra BWM and NovoSorb BTM in pig skin wounds. EG7 PTK-UR foams performed as well as Integra in terms of wound closure and tissue repair. Though similar rates of wound closure were observed between Integra and EG7 PTK-UR treatments, significantly higher blood perfusion was seen in wounds treated with EG7 PTK-UR, indicating that vascularization is an especially strong feature of EG7 PTK-UR scaffolds (Figure 7C). A slower degradation profile of NovoSorb was inferred by the presence of a relatively large proportion of scaffold material in the wound after 31 days (Figure 7D), which could potentially cause a prolonged wound inflammatory response. Similarly, long-term persistence of NovoSorb scaffold remnants has been reported in human skin biopsies take 12 months post implantation (*98*). This may rationalize the relative upregulation of pro-inflammatory cytokines such as IL1β (3.1-fold), TNFα (2.1-fold), NOS2 (4.5-fold), CXCL8 (12-fold), and CSF2 (5.9-fold) observed in NovoSorb treated wounds compared to EG7 PTK-UR treatments (Figure 8D). In comparison to architecturally and synthetically analogous NovoSorb foams, EG7 PTK-UR foams explanted from wounds at day 31 had significantly upregulated expression of anti-inflammatory interleukins IL13 (2.8-fold) and IL33 (3.2-fold), which are known to induce a tissue repair phenotype in macrophages (*99, 100*). Surprisingly, we observed significantly lower levels of arginase-1 (10-fold) expression in EG7 PTK-UR wounds relative to NovoSorb, a marker often used to characterize the anti-inflammatory M2 macrophage phenotype. Arg-1 IHC of wound tissue revealed a decrease but not the absence of Arg-1 expressing macrophages in EG7 PTK-UR treated wounds relatively to NovoSorb. Intense Arg-1 staining was, however, observed colocalized with FBGCs surrounding NovoSorb scaffold remnants. Despite the importance of arginase-1 metabolism in macrophage polarization, elevated levels of arginase-1 have been reported in ischemic porcine wounds (*101*) and in human venous leg ulcers (*101*). In this context, high arginase activity has been linked to fibrosis (*102*), suggesting disparate biological ramifications of Arg- 1expressing macrophages. Relative abundance of TGFβ3 expression relative to TGFβ1 (Figure 8C) has been shown to promote scarless healing in mice compared to excess ECM deposition and fibrosis seen with TGFβ1 upregulation (*103*).

This study has limitations that should be discussed. First, testing the regenerative capacity of PTK-UR in acute wounds created on healthy pigs gives important but still limited insight into performance in chronic human skin wounds. Chronic wounds are associated with greater ischemia, in many cases infection, and potentially higher levels of oxidative stress that may over-accelerate the degradation of EG7 PTK-UR scaffolds. Additional tuning of mechanical properties to compensate for accelerated degradation maybe be required in order to effectively treat chronic wounds. Another limitation is the small wound size used in the study. Reepithelization of larger wounds with EG7 PTK treatment alone will be challenging due to limited migratory capacity of keratinocytes on granulation tissue. The successful use of EG7 PTK-UR scaffolds for larger skin wounds would most likely be paired with the application of split-thickness skin graft or autologous keratinocyte transplant (*104*).

In summary, we have developed a novel EG7 PTK-UR foam dressing that facilitates bulk tissue-scaffold integration, robust cellular and vascular infiltration, and wound re-surfacing. These implants induce a moderate inflammatory response that effectively transitions to a pro-healing phenotype in vivo, in addition to promoting ECM deposition and remodeling. When tested against clinically approved materials in healing of porcine excisional wounds, EG7 PTK-UR performed at par with the gold standard, Integra BWM, and significantly outperformed the synthetic polyester-based foam NovoSorb.

## Acknowledgments

The authors acknowledge the Translational Pathology Shared Resource supported by National Cancer Institute (NCI)/NIH Cancer Center Support Grant 2P30 CA068485-14, Vanderbilt Mouse Metabolic Phenotyping Center Grant 5U24DK059637-13 and Digital Histology Shared Resources at Vanderbilt. Funding was provided by the National Institutes of Health (NIH) through R01EB019409 and the Department of Veterans Affairs. We confirm that there are no known conflicts of interest associated with this publication and that there has been no significant financial support for these efforts that could have influenced their outcome.

## Supplemental Figures

**Figure S1:**
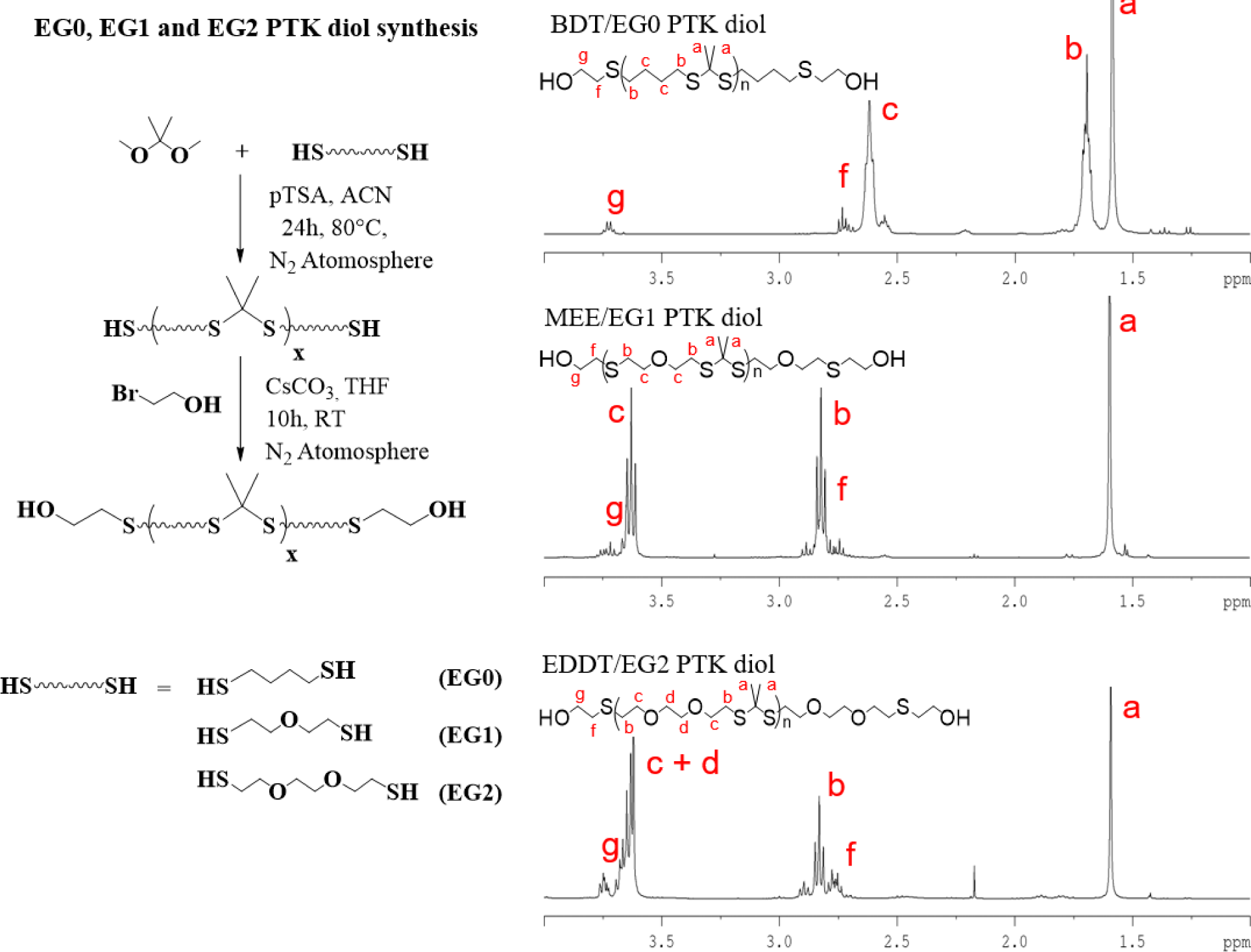
Synthesis and H NMR characterization of EG0, EG1 and EG2 PTK diols. (A) **Synthesis** scheme for condensation polymerization of BDT (EG0), MEE (EG1) and EDDT (EG2) dithiol monomers to obtain PTK dithiol polymers and their functionalization with bromoethanol to synthesize PTK diol polymer. On the right, ^1^ H NMR spectra of BDT PTK diol, MEE PTK diol and EDDT PTK diol showing accurate chemical composition.

**Figure S2:**
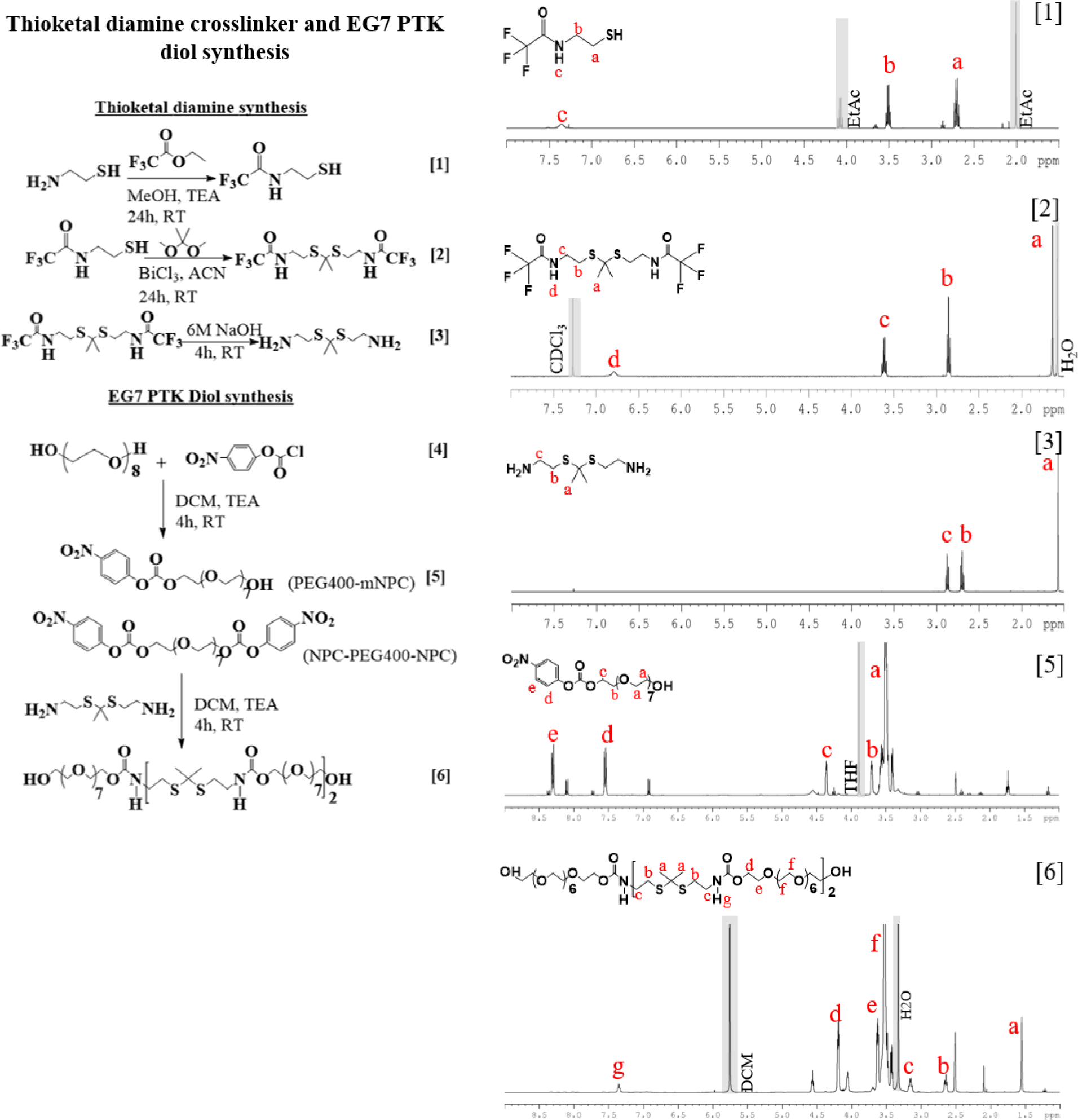
Synthesis and ^1^H NMR characterization of EG7 PTK diol. (A) ROS degradable thioketal diamine crosslinker and EG7 PTK diol through the reaction of TK diamine and amine reactive PEGmNPC/NPC-PEG-NPC. On the right, ^1^H-NMR spectroscopy of compounds 1, 2, 3, 5 and 6 are shown.

**Figure S3:**
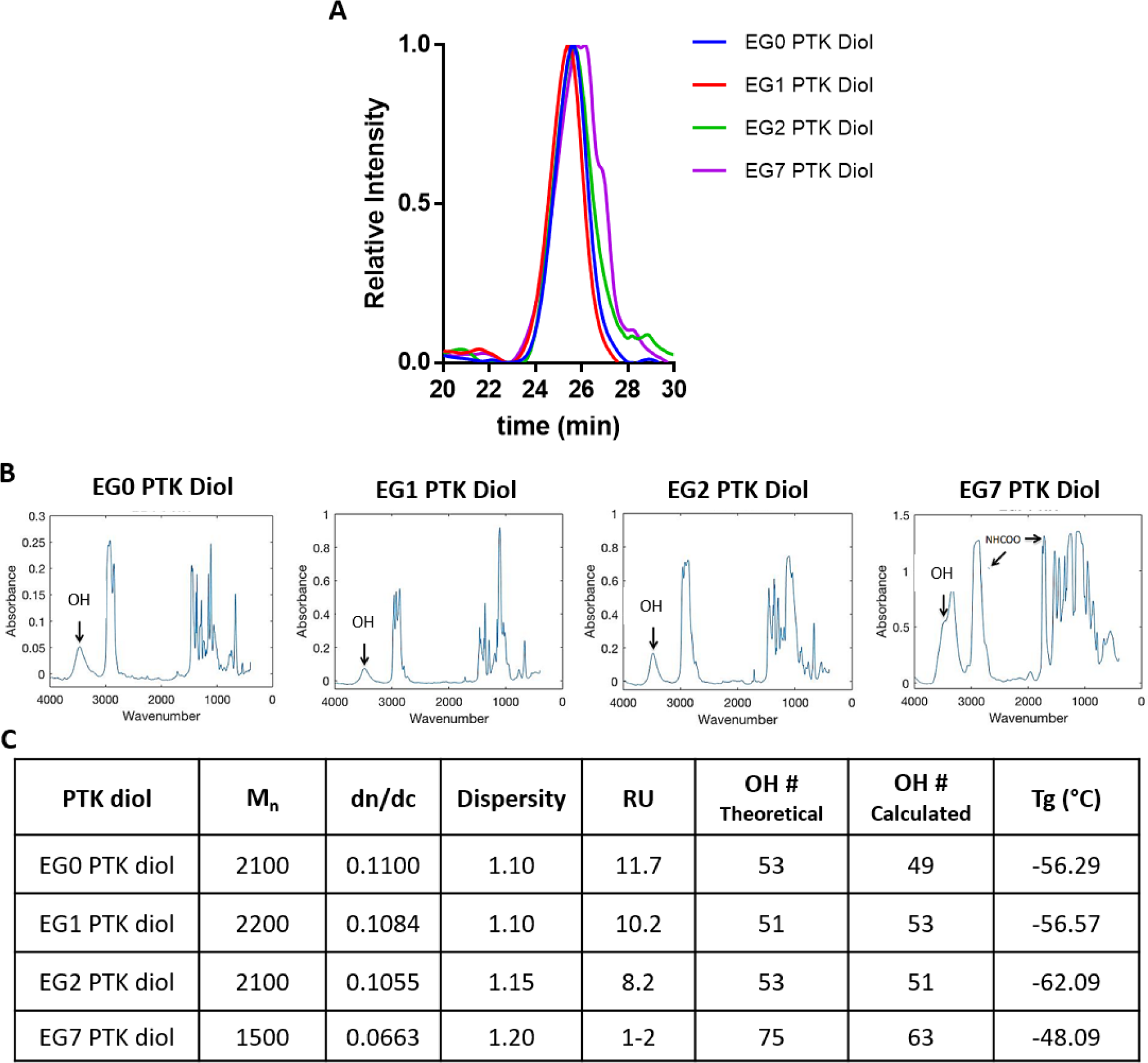
Characterization of PTK diols. (A) GPC elutograms for EG0, EG1, EG2, and EG7 polymers. (B) FTIR spectra of hydroxyl terminated PTK polymers with hydroxyl absorbance peak seen at 3400 cm-1 (black arrow). (C) Table showing number average molecular weight (Mn), dispersity index, thioketal repeating unit (RU), hydroxyl number (OH#, calculated through 19F NMR), and glass transition temperate (Tg) of EG0, EG1, EG2, and EG7 PTK diols

**Figure S4:**
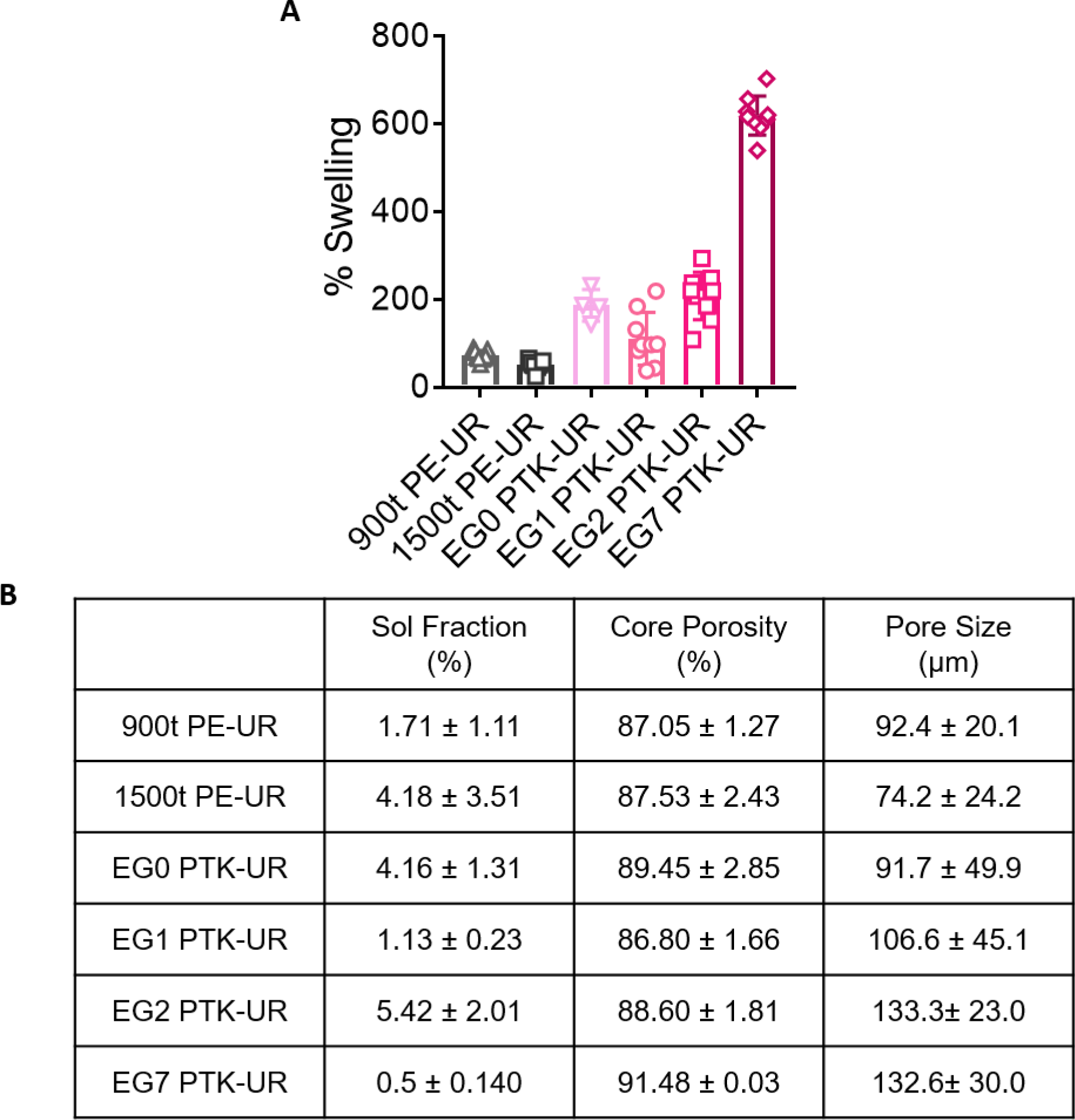
(A) Swell ratio of PE and PTK based urethane foams hydrated in PBS, (B) Physical properties of PE-UR and PTK-UR scaffolds.

**Figure S5:**
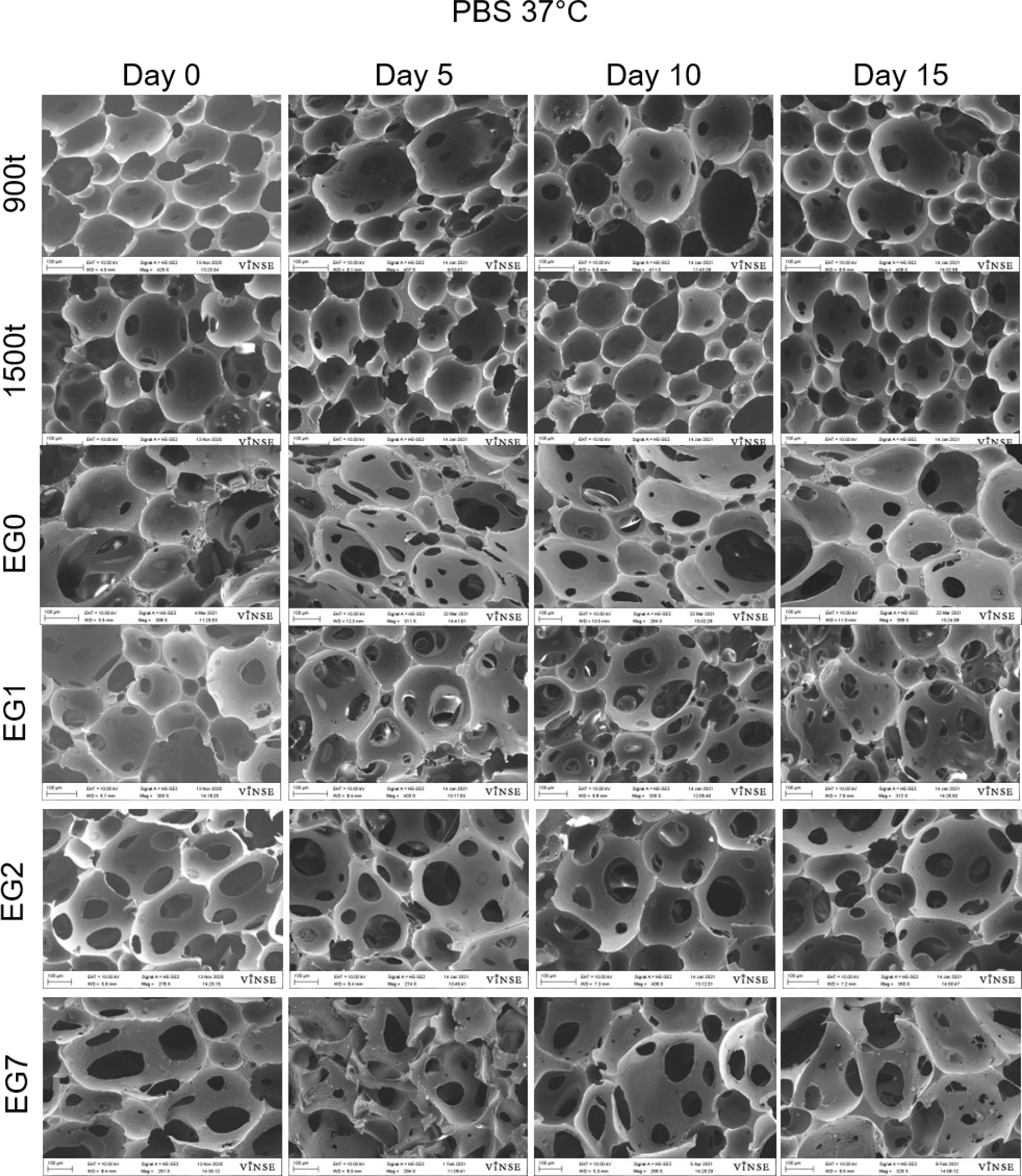
Representative SEM images of in vitro PTK and PE-UR scaffold degradation incubated in hydrolytic media (PBS) for 15 days.

**Figure S6:**
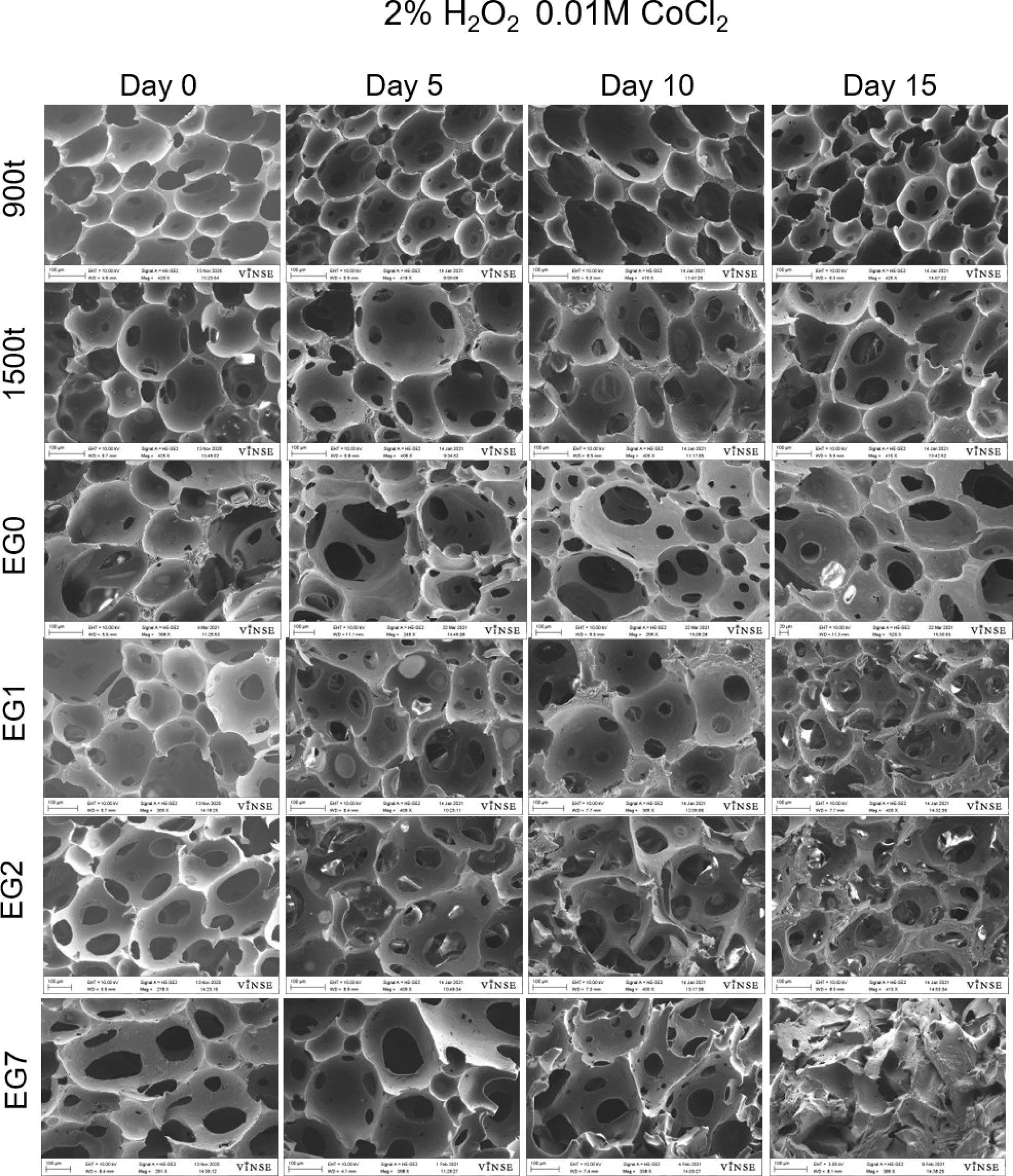
Representative SEM images of in vitro PTK and PE-UR scaffold degradation incubated in oxidative media simulated by 2% H_2_O_2_ 0.01M CoCl_2_ for 15 days.

**Figure S7:**
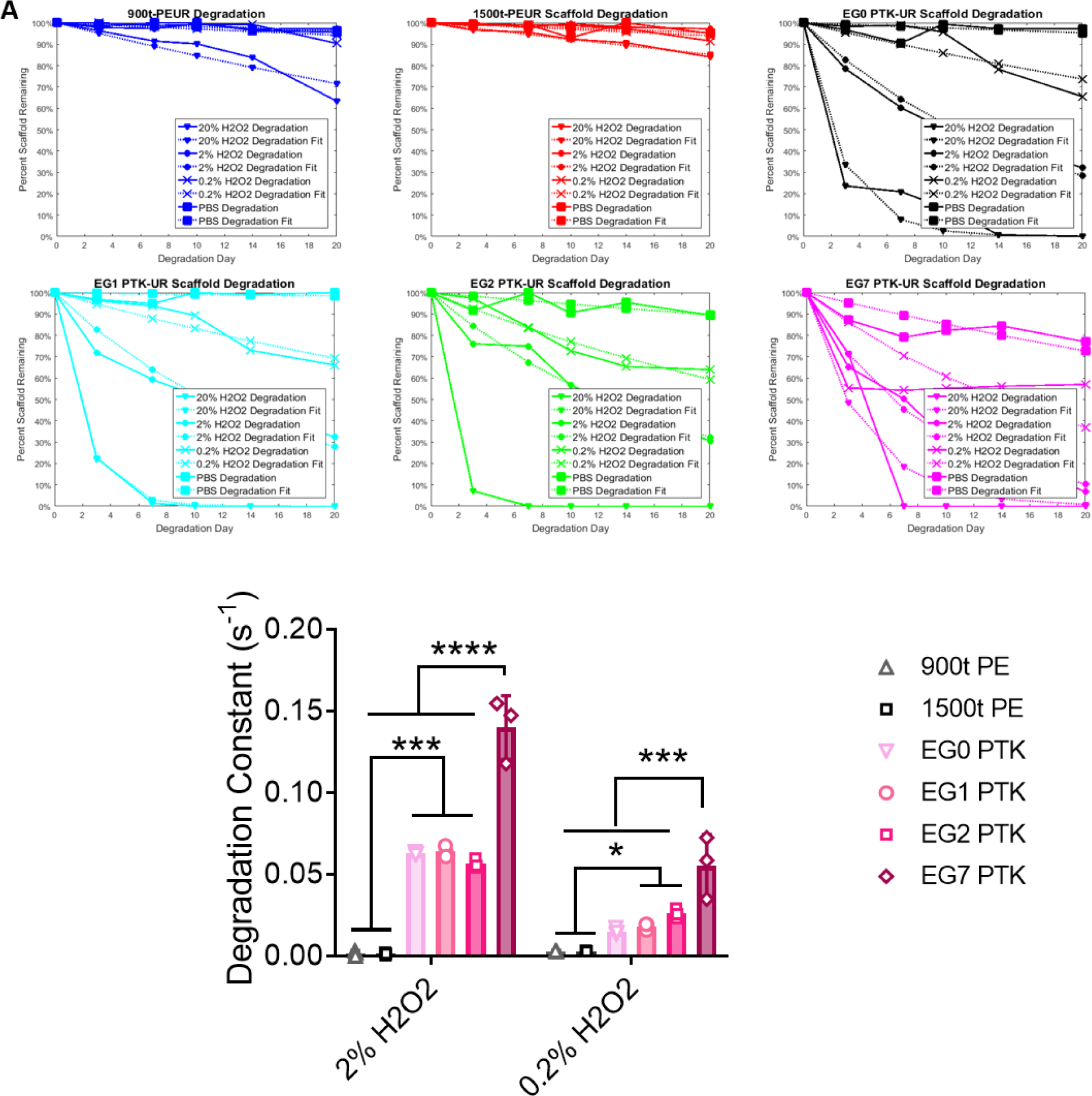
(A) In vitro degradation kinetics of PTK-UR scaffolds showing dose dependent degradation in oxidative media and relative stability in hydrolytic media over 30 days. Dashed lines represent MATLAB model-generated curves for degradation kinetics. (B) Degradation constants derived from the experimental data shown above.

**Figure S8:**
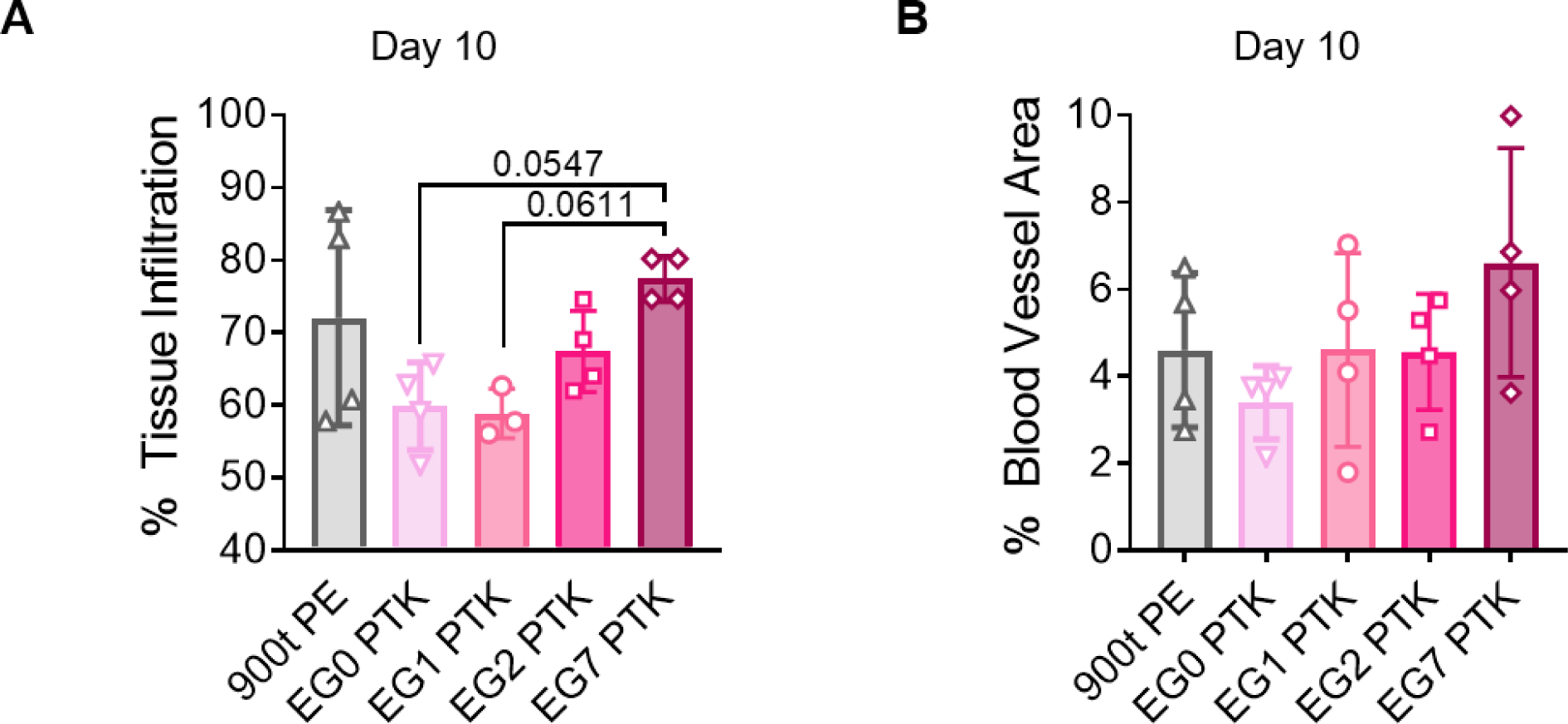
Quantification of (A) tissue infiltration (B) blood vessel area within scaffold infiltrating tissue 10 days post-wounding.

**Figure S9:**
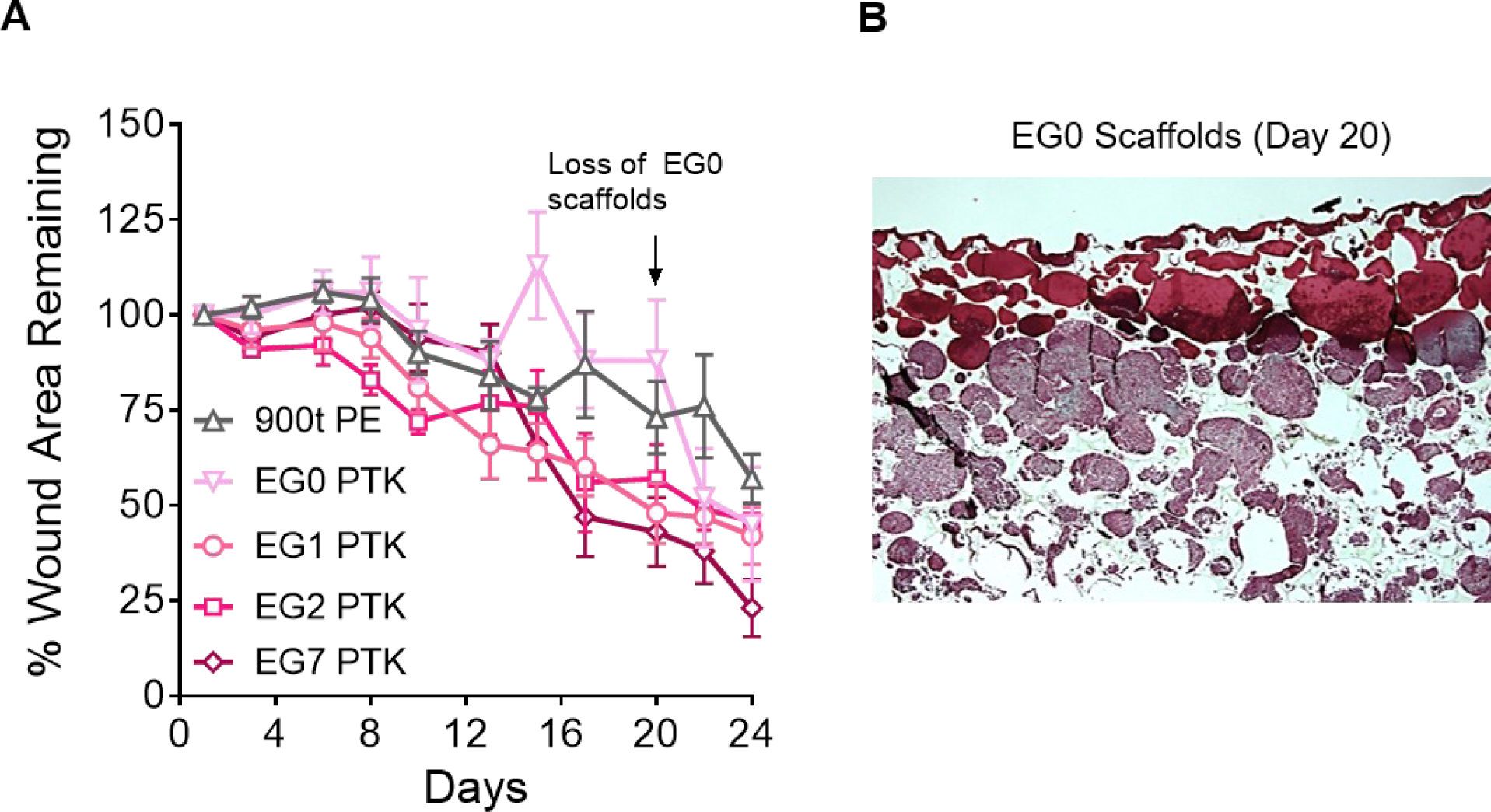
(A) Quantification of wound area remaining open over a period of 24 days of scaffold treated porcine skin wounds. (B) Representative trichrome image showing tissue infiltration within EG0 PTK- UR scaffold pores 20 days post implantation. Scaffold were extruded from the wound post application.

**Figure S10:**
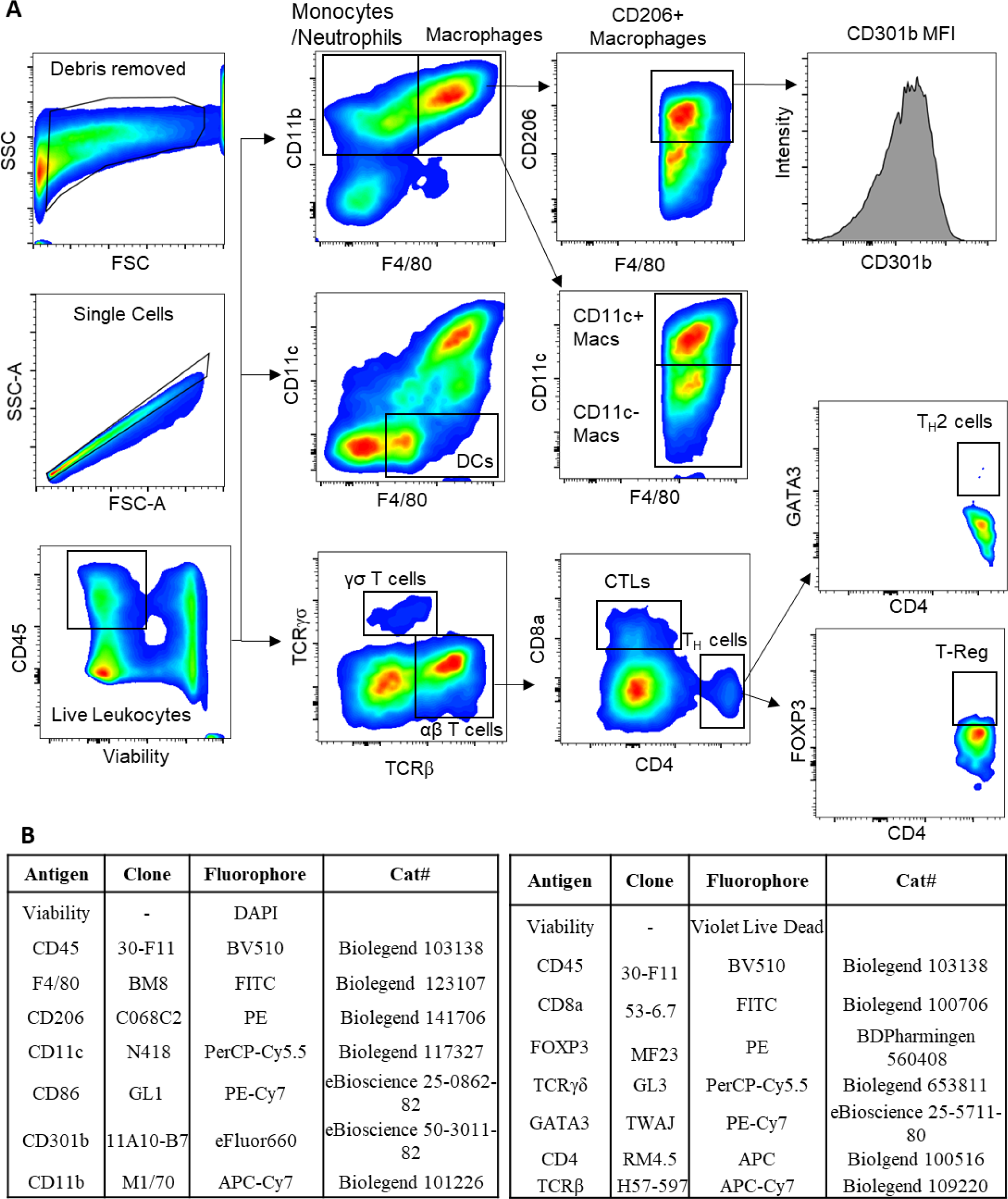
(A) Gating strategies for myeloid and lymphoid populations. Cells were gated on SSC and FSC, followed by doublet discrimination to isolate the single cell population based of FSC-A and FSC-H. Cells were gated on live immune cells (DAPI+CD45+) or (Viability dye+CD45+) prior to phenotypic specific gating (B) Antibody clones used in flow cytometry and immunophenotyping of myeloid and lymphoid cells

**Figure S11:**
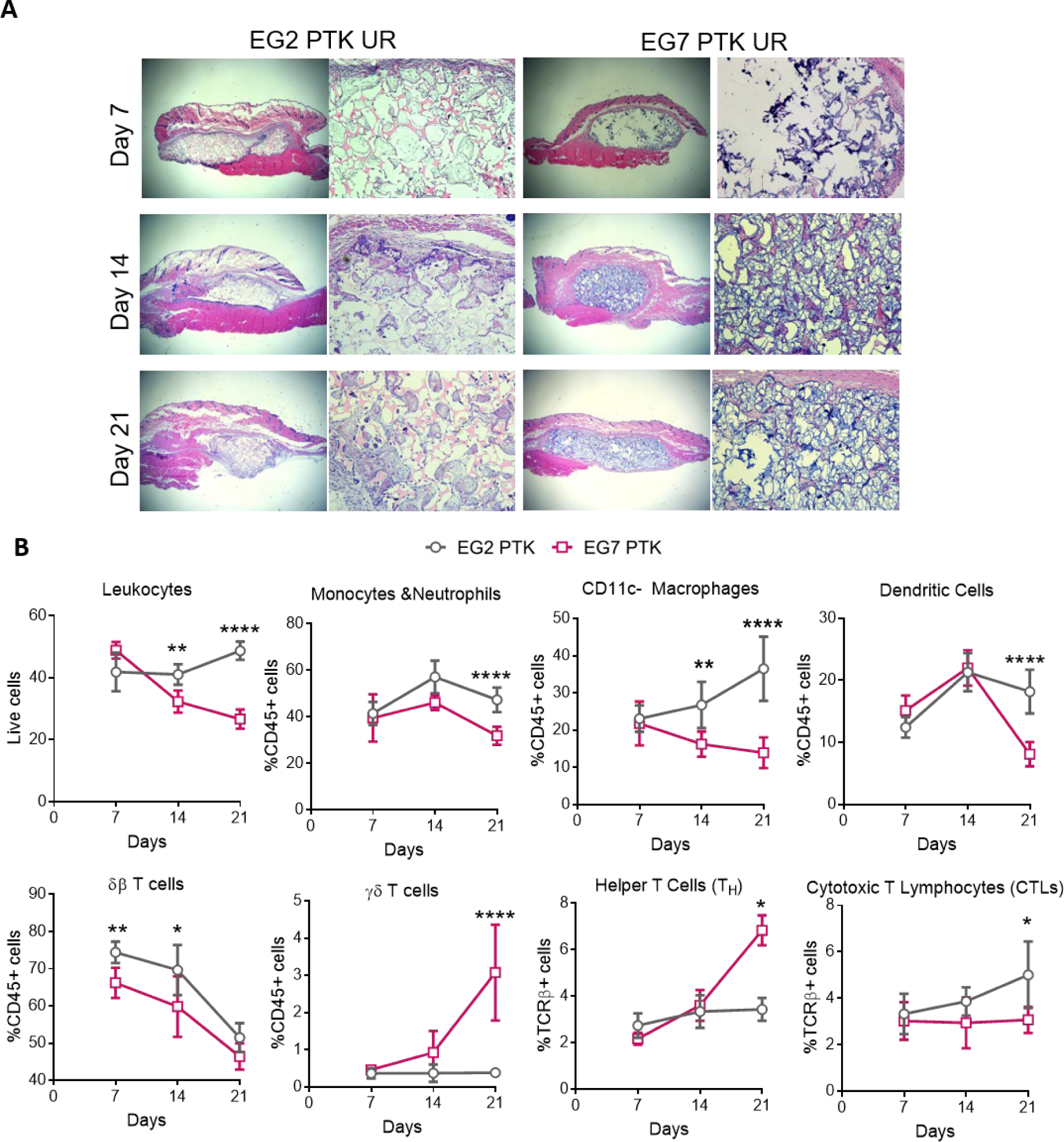
(A) 6mm x 1.5mm EG2 and EG7 PTK-UR scaffolds implanted in subcutaneous pockets of C57BL/6 mice and explanted after 7, 14 and 21 days. Representative H&E images showing tissue infiltration and immune cell recruitment into scaffold pores. (B) Change in leukocyte sub-populations in EG2 PTK (gray) and EG7 PTK (pink) cellular infiltrates measured 7-, 14- and 21-days post implantation.

**Figure S12:**
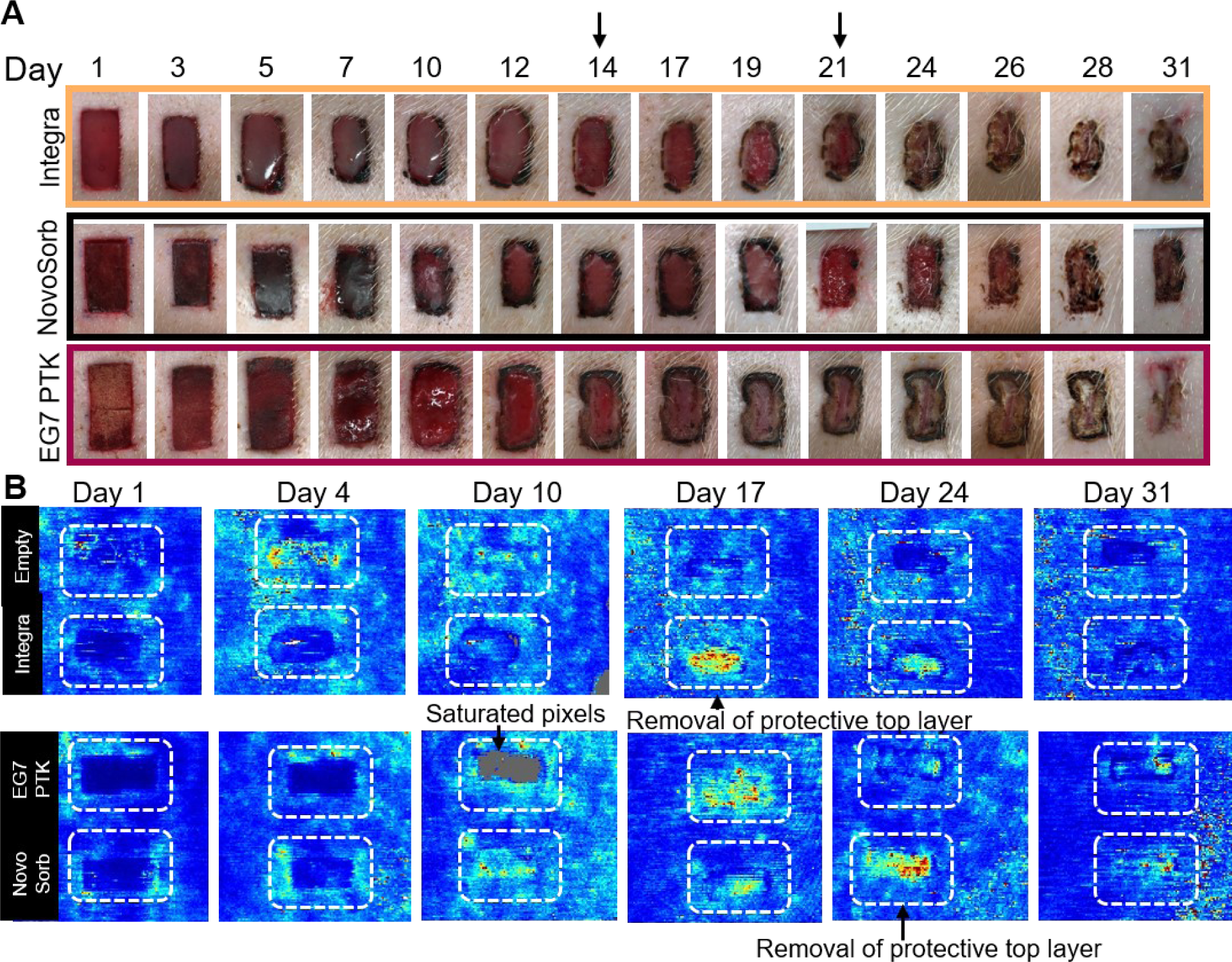
(A) Photographic images of porcine skin wounds treated with Integra BWM, NovoSorb BTM and EG7 PTK-UR scaffolds showing remaining wound area and scaffold resolution. Arrows indicating removal of Integra protective silicone layer on day 14 and BTM polyurethane layer on day 21. (B) Representative LDPI flux images showing blood perfusion within the wound. Removal of protective top layer results in re-wounding of Integra and NovoSorb treated wounds

**Figure S13:**
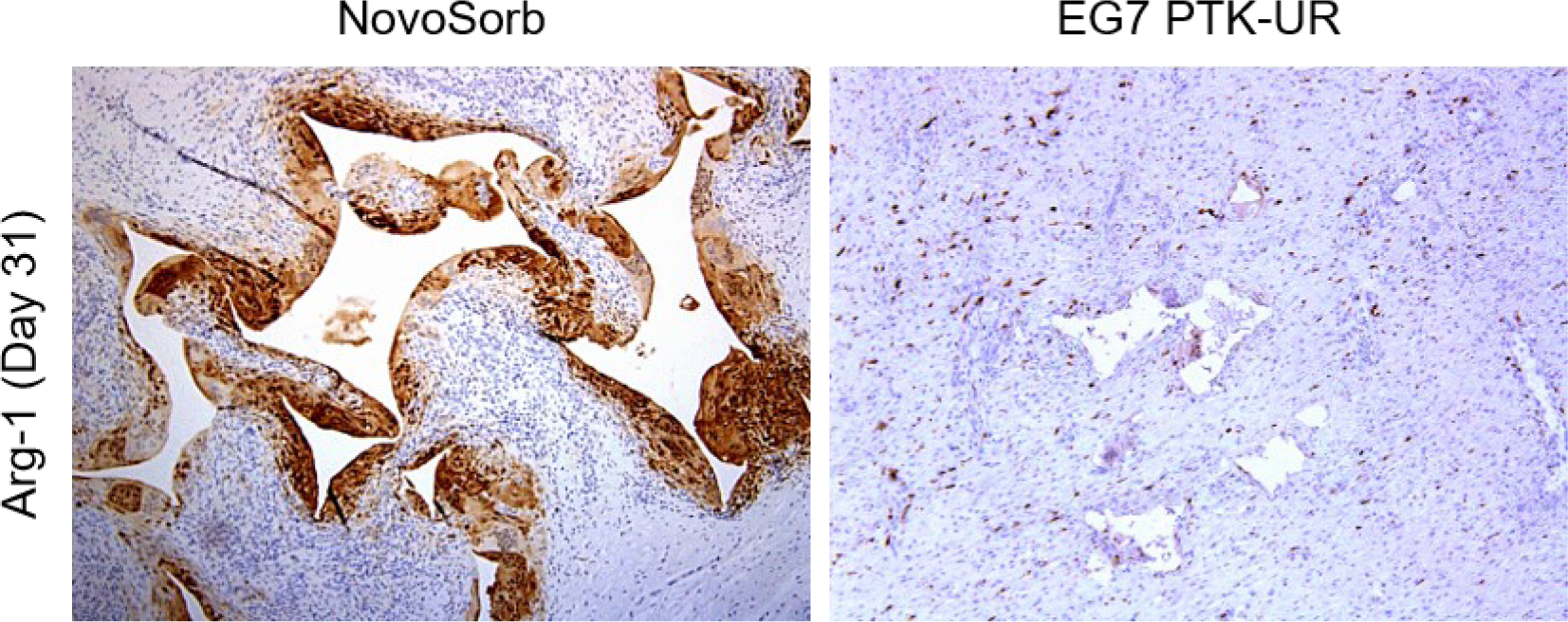
Arginase-1 IHC of NovoSorb and EG7 treated wounds.

